# Evaluating a harvest control rule to improve the sustainability of Japanese fisheries

**DOI:** 10.1101/2020.07.16.207282

**Authors:** Hiroshi Okamura, Momoko Ichinokawa, Ray Hilborn

## Abstract

Fisheries management in Japan is currently at a turning point. MSY based reference points have historically been rejected because of impacts on the fishing industry that would result from their adoption. We propose and evaluate a new harvest control rule (HCR) that uses the biological reference points based on sustainable yield from the stochastic hockey-stick stock recruitment relationship. Management strategy evaluation simulations conditioned on data from Japanese stocks demonstrate that the new HCR avoided recruitment overfishing while providing stable and near maximum catch. The new HCR outperformed Japan’s traditional HCR in terms of conservation, and it outperformed an alternative HCR which is widely used around the world in terms of initial catch reduction and future catch variation. For forecasting and hindcasting simulations, the new HCR showed considerable improvements over traditional HCRs in terms of biomass and catch. This new management procedure can improve the current and future status of many overfished stocks in Japan as well as increase economic efficiency and better protect ecosystems.

## 1. Introduction

Harvest control rules (HCRs) and management strategy evaluation (MSE) play an important role in global contemporary fisheries management (Smith 1994; De la Mare 1998; Butterworth and Punt 1999; Deroba and Bence 2008; Froese et al. 2011; Hillary et al. 2016; Punt et al. 2016, 2017). Because of the inevitable trade-offs between long- and short-term objectives in fisheries management and the large uncertainties inherent in marine ecosystems, selecting an HCR is a complex task. MSE is a powerful approach for comparing alternative HCRs from a range of choices and achieving management targets in the presence of large uncertainties.

Scientific and non-scientific experts in Japan have rejected biological reference points (BRPs) based on the maximum sustainable yield (MSY) as management targets for many years because of uncertainty about how to estimate the MSY reference points. Purportedly, these uncertainties arise from ecosystem complexity resulting from regime shift and multispecies interactions (Kidokoro et al. 2010; Matsuda and Abrams 2006). There is evidence that the environment more strongly affects recruitment (R) than spawning biomass (SB) in many stocks (Sakuramoto 2005; Szuwalski et al. 2015). By contrast, Ichinokawa et al. (2017) demonstrated that the primary fish stocks in Japan have relatively low R variation: the variability around the SR relationships (CV_R_) was 0.39 for Japan (Ichinokawa et al. 2017) compared to 0.74, globally (Thorson et al. 2014). In addition, R and SB were positively related over the observed range of stock sizes in 64% for Japan (Kurota et al. 2020) compared to 39% for the world (Szuwalski et al. 2015). Thus, stock-recruitment (SR) relationships in major Japanese fisheries stocks have been well estimated compared to those in worldwide marine fisheries (Szuwalski et al. 2015). Costello et al. (2016) and Ichinokawa et al. (2017) argued that Japan’s fisheries had low stock levels and could increase yields and recover quickly by reducing fishing intensity to appropriate levels. Thus, Japanese national fisheries badly need a new target-based management approach that is capable of addressing large uncertainties such as climate change.

There is circumstantial evidence that reducing fishing pressure in Japan leads to quick recovery in declining fish stocks. The Great East Japan Earthquake that struck Japan on March 11, 2011 unleashed giant tsunamis that destroyed seaside cities and damaged the Fukushima Daiichi nuclear power plant. Although it was recently estimated that radioactive contamination from the nuclear plant accident did not seriously harm marine fish (Okamura et al. 2016), many fisheries still remained closed or restricted in Fukushima. During this period, some species of fish off the coast of Fukushima have become increasingly abundant, probably because of significantly reduced fishing pressures (Narimatsu et al. 2017; Shibata et al. 2017). Reducing fishing pressures in other areas has had similar positive effects on abundance (Neubauer et al. 2013; Hilborn et al. 2020); in other words, eliminating overfishing could rapidly lead to higher fish abundance, and more stable catches.

The Japanese government has not fully implemented modern fisheries management based on BRPs into the Japanese fisheries system (Matsuda et al. 2010; Makino 2011), and MSY-based targets have been actively avoided. Rather, maintaining the status quo for stocks that have recovered from low levels has been the only agreed-upon management target for data-rich stocks (Watanabe 2018). However, with analytical and empirical evidence supporting the effectiveness of reducing fisheries efforts (Fernandes and Cook 2013; Neubauer et al. 2013; Ichinokawa et al. 2017; Shibata et al. 2017; Hilborn et al. 2020), the Japanese government is launching a new fisheries management system that includes explicit MSY-based management targets for each stock (https://www.spf.org/en/spfnews/information/20190830.html). Now, a variety of movements have begun to demand the improvements in Japan’s fisheries management and the methods that bridge the gap between MSY theory and practice and are robust to uncertainties.

Because it has not been necessary to estimate the MSY, the commonly used age-structured population dynamics model in Japan has been a virtual population analysis, which does not consider SR relationships (Hiramatsu 2018). Therefore, in many domestic stock assessments, the SR relationships were not explicitly or formally established. Based on current stock assessment data in Japan, we use the approach of Ichinokawa et al. (2017) to estimate BRPs at MSY.

As a component of emerging fisheries management systems in Japan, we propose a new HCR based on BRPs at MSY which is an approach that should be suitable for Japanese fisheries stocks with rich stock assessment data. In the new HCR, BRPs are estimated by a simulation assuming a stochastic hockey-stick (HS) SR curve to take both the importance of spawning stock and natural variability into account (Ichinokawa et al. 2017). Estimating the SR curve and BRPs are usually conducted in the stock assessment and used as input data into the MSE simulation. However, we estimate them by fitting the SR curve to the SR time series data with stochastic errors generated from the operation model in our MSE simulation to accurately propagate the estimation uncertainty of the SR relationship (the technical detail is given in the Method section). A merit of using a stochastic HS SR curve, whatever the true SR curve is, is discussed later. Using extensive simulation tests with various sources of uncertainty, we investigate whether the new HCR outperforms the status quo HCR that has been used in Japan and the HCR used typically around the world (40–10 rule; Deroba and Bence 2008). We quantify the impact of using the new HCR on Japanese fisheries by summarizing the simulation results. The remainder of the paper is structured as follows. Section 2 describes the methods, including the HCRs and the simulation specifications. Section 3 presents the results from the simulations and the real data analysis. Section 4 discusses the interpretation and the indication from the results and the future direction of Japan’s fisheries management.

## 2. Methods

### 2–1. Estimation of BRPs using a stochastic HS curve

BRPs, such as SB at MSY (SB_msy_) and fishing rate at MSY (*F*_msy_), had not been estimated for most Japanese stocks until recently because of uncertainty in the SR relationships and a distrust of density dependence and the MSY concept (e.g. Sakuramoto 2005). In Japanese fisheries, population abundance has usually been estimated via virtual population analysis without assuming any specific SR relationship (Ichinokawa et al. 2017; Okamura et al. 2017). Future projections were conducted by resampling the recruits per spawner. Because this approach has no limit on potential recruitment, this approach sometimes produced a large and unrealistic future biomass. In these cases, the upper limit of R was often imposed in an ad hoc manner (Yoda et al. 2019). This means that researchers implicitly assumed a HS curve as a SR relationship. Thus, we adopt a stochastic HS curve including the R variation as the basic SR relationship for stocks in Japan (Ichinokawa et al. 2017).

A stochastic HS curve has the following characteristics:

1. The deterministic value of recruitment declines linearly below a spawning biomass threshold and is constant above the threshold, which is easily understood by non-scientific experts.
2. It does not lead to extremely large (or small) values of SB_msy_ being extrapolated beyond the historical SB time series (Ichinokawa et al. 2017), by defining the minimum observed SB for the flat SR curve and the maximum observed SB for the linear SR curve as the threshold (Appendix S1). This circumvents estimating extremely large or small reproductive potential (aka steepness), which we come across using well-known SR functions such as the Beverton-Holt and Ricker functions (Hilborn and Walters 1992; Punt et al. 2014), though the estimated SB_msy_ is not always within the range of historical SBs (In a stochastic environments, SB_msy_ using the HS curve is generally larger than the threshold (Appendix S2)).
3. It is robust against uncertainty because *F*_msy_ becomes lower and SB_msy_ becomes higher as R variation increases (Appendix S2). This is consistent with a precautionary approach.
4. It has a wide range of SB generating sustainable yield close to MSY as the form of the curve is similar to the Beverton-Holt curve with a plateau for high SB (Hilborn 2010; Punt et al. 2014). These HS characteristics permit stakeholders who may face trade-offs when setting management objectives to accept BRPs in a smooth and timely fashion.

We can consider the HS curve as a proxy for an asymptotically flat-topped SR curve such as a Beverton-Holt curve. HS provides the MSY correctly when the true SR relationship is an HS with a break point within the observed SBs, which prevents controversies such as ‘Is 30 or 40% SB_0_ better?’. Having a constant R that exceeds a threshold is easily understood and acceptable by many fish stock assessment researchers and non-scientific experts, alike.

However, the simplicity of HS masks some of its complication, which are caused by indifferentiability at the break point of the curve (Mesnil and Rochet 2010). Unless the growth effect and variation in R are considered, the break point of the deterministic HS model corresponds to SB_msy_ and SB_crash_ (where the population goes to extinction when harvesting beyond *F*_msy_) at the same time. In terms of stochastic variation in R, SB_msy_ tends to be found at a higher SB than the break point, because SB at the break point produces a reduced R on average (Ichinokawa et al. 2017; Appendix S2). In terms of growth, SB_msy_ becomes the break point or SB fished at *F*_max_ (SB_max_), which is the fishing rate that gives the maximum yield per recruit (Quinn and Deriso 1999), depending on the location of SB_max_. Taking the growth effect and stochastic variation in R into consideration, we define the maximum value of the long-term average yield as (stochastic) MSY (Ichinokawa et al. 2017) and refer to SB producing MSY under a stochastic environment as SB_msy_.

SB_msy_ is our target SB, SB_tar_. Because modern HCRs often utilize the form in Fig. 1 (Deroba and Bence 2008; Froese et al. 2011; Thorson et al. 2015), we have a further need for SB_lim_ and SB_ban_ that are other BRPs for fisheries management based on HCRs. SB_lim_ is a threshold for ensuring secure sustainable yields, and SB_ban_ is a threshold for avoiding irreversible outcomes such as Allee effects (Perälä and Kuparinen 2017). When SB drops below SB_lim_, the fishing mortality rates are reduced and when SB is below SB_ban_, fisheries are closed.

**Figure 1.**
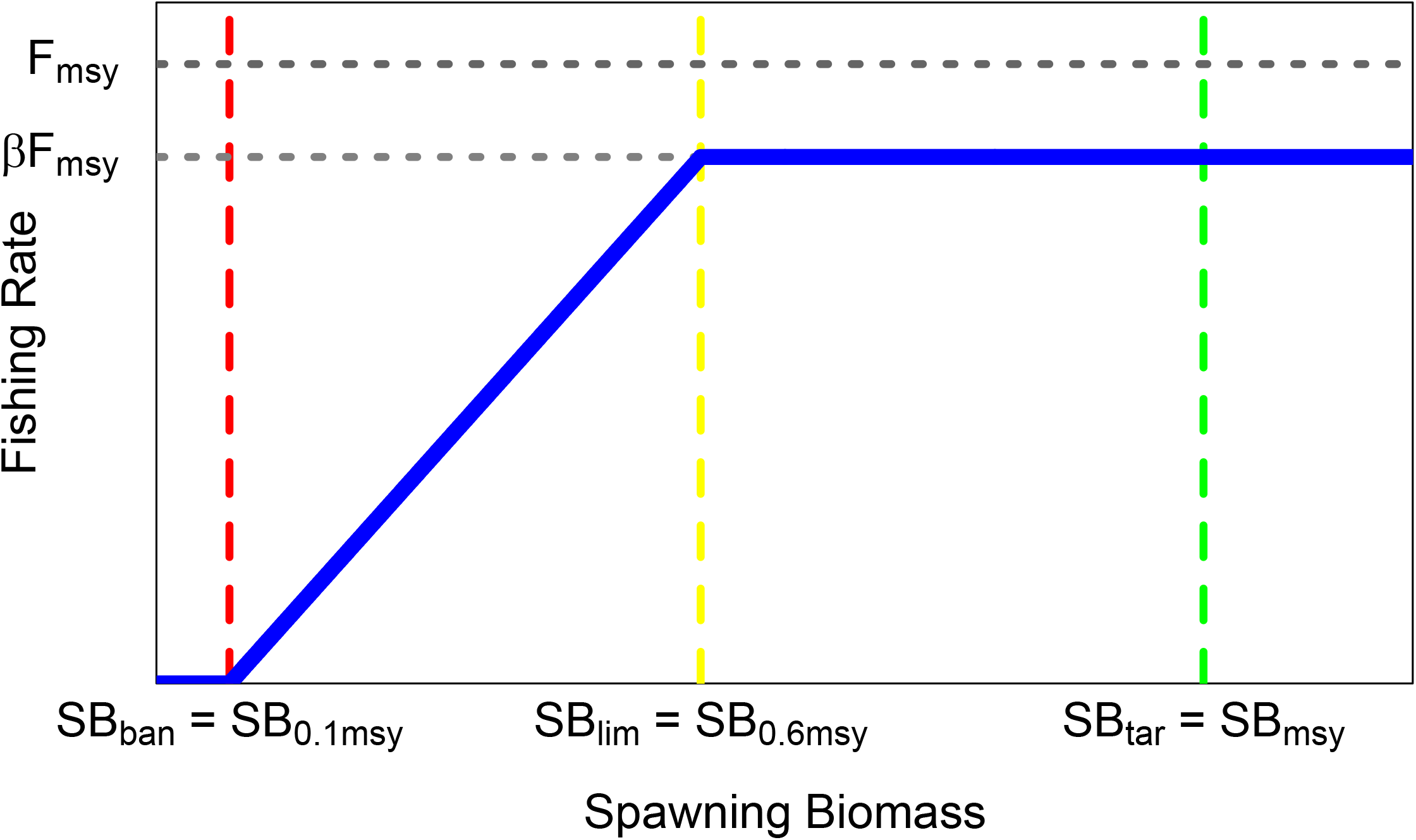
New harvest control for Japanese data-rich fisheries.

When fitting HS to the SR data, we use the method of least absolute deviations for log R data to take large fluctuations in R into account and avoid excessive impacts of strong cohorts. We then estimate the SR parameters *a* and *b* by minimizing

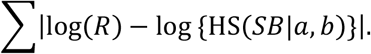

We then fit the first-order autoregressive model, *r*_*t*+1_ = *ρr*_*t*_ + *ε* where *ε* ~ N(0, *σ*^2^), to the residuals in log R in year *t*, *r*_*t*_ = log(*R*_*t*_) − log {HS(*SB*_*t*_|*a*, *b*)}. For the autoregressive model of the residuals, when the AIC of the model without autocorrelation is smaller than that of the model with autocorrelation, we set the autocorrelation to zero. When we estimate the MSY-related parameters and conduct future projection under each HCR, we attach the estimated autocorrelated errors to R in the simulation (Appendix S3).

### 2–2. HCRs

Acceptable biological catch (ABC) in year *t* is calculated by ABC_*t*_ = *F*_*t*_ × *B*_*t*_ according to the following three HCRs, where *F*_t_ is the fishing rate and *B*_t_ is the total biomass in year *t*. Because ABC in Japan is usually determined by the two years ago biomass estimates, we consider two cases, one with no time lag (the case determined by one year ago biomass estimates) and one with a time lag of one year (the case determined by two years ago biomass estimates), to examine the impact of time lag on HCR management. The mathematical details are given in Appendix S3.

#### 2–2–1. Japan’s traditional HCR

In many Japanese fisheries stock assessments, the SR curve is not formally used. Instead, SB_lim_ is sometimes estimated by the ‘sb’ method, which determines the threshold by the intersection between the 90%-ile points of R and RPS (Serebryakov 1991; Myers et al. 1994). When SB_lim_ is determined, the *F* value is provided by

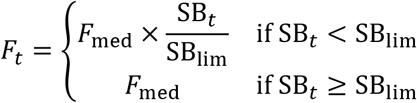

where *F*_med_ is the *F* value that maintains the median of SPR (SB per R) without stochastic simulation (i.e., deterministically) (Cadima 2003). While SB_lim_ is updated every five years, *F*_med_ is calculated every year because the current ABC control rule in Japan does so. We refer to this HCR as simply ‘Status Quo’. In Status Quo HCR, SB_ban_ is set to zero, as currently many stocks do not have SB_ban_.

#### 2–2–2. The 40–10 HCR

40–10 HCR has been accepted by the Pacific Fisheries Management Council, and some MSE simulations have demonstrated that it has good performance (Thorson et al. 2015; Benson et al. 2016). In 40–10 HCR, SB_lim_ is 0.4×SB_0_ and SB_ban_ is 0.1×SB_0_ where SB_0_ is the equilibrium value for SB in an unfished population. The *F* value is then provided by

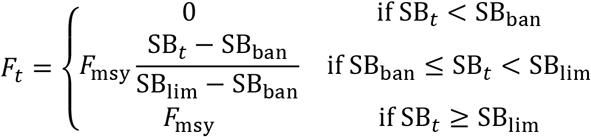

Since we do not have *F*_msy_ and SB_0_ from the formal stock assessments, we estimated them from the simulation using the stochastic HS curve. While *F*_msy_, SB_lim_, and SB_ban_ (and SB_0_) are updated every five years, SB_*t*_, B_*t*_, and ABC are updated every year. We call this HCR ‘40–10’.

#### 2–2–3. The new HCR

In the new HCR, SB_tar_ = SB_msy_, SB_lim_ is SB_0.6msy_ which is SB (< SB_msy_) producing 60% of MSY under a constant fishing rate (*F*), and SB_ban_ is SB_0.1msy_, which is SB (< SB_msy_) producing 10% of MSY under a constant *F*. Because HS does not have a stable equilibrium below the break point, SB_0.6msy_ and SB_0.1msy_ do not always exist. We therefore use the 20 times generation time forward simulation under a constant *F* to obtain approximate SB_0.6msy_ and SB_0.1msy_. Preliminary analyses using Japanese fisheries stock data have indicated that increasing the generation time produced almost the same BRPs.

The *F* value is then provided by

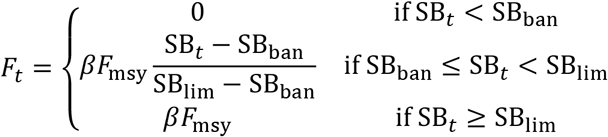

where *F*_msy_, SB_lim_, and SB_ban_ are estimated from the simulation using the stochastic HS curve. The adjustment parameter *β* is fixed at 1 or 0.8 in this paper. While *F*_msy_, SB_lim_, and SB_ban_ are updated every five years, SB_*t*_, B_*t*_, and ABC are updated every year. We call this HCR with *β* = 1 ‘NEW’ and this HCR with *β* = 0.8 ‘NEW2’ (Fig. 1).

### 2–3. Simulation

#### 2–3–1. Reference scenario sets

The simulated data are generated by using the delay-difference model (Hilborn and Walters 1992) with the HS SR relationship. The delay difference model is an intermediate model between age-aggregated surplus production models and age-structured population dynamics models, and explicitly expresses biological processes such as survival, growth, and recruitment (Appendix S3).

To conduct the stochastic simulation incorporating the information from the real stock assessment data, we carry out the following ‘conditioning’ on the information from the stock assessments. Based upon a deterministic *F*_msy_ (= MSY/*B*_msy_) and *b*_ratio_ (= b/SB_0_ where *b* is the break point of HS) obtained by fitting HS to 26 stocks with SR data in Ichinokawa et al. (2017) (Fig. 2), we determine the ranges of the natural survival rate (*S*), *F*_msy_, and *b*_ratio_: *S* ∈ {0.37, 0.67, 0.82} (these values correspond to natural mortality coefficients *M* = 1.0, 0.4, and 0.2, respectively), *F*_msy_ ∈ {0.10, 0.25, 0.40}, and *b*_ratio_ ∈ {0.10,0.25,0.40}. Although this produces 3×3×3=27 combinations in total, all combinations are not always realistic for Japanese fisheries due to correlations between biological parameters (Fig. 2). To select the realistic combinations, we first fit the trivariate normal distribution to the combinations of (*S*, *F*_msy_, *b*_ratio_) for 26 Japanese fish stocks in Ichinokawa et al. (2017) (Fig. 2), then calculate the probability densities (or likelihoods) for 27 combinations from the fitted trivariate normal distribution, and eliminate the implausible combinations with the standardized probability < 0.001. We thus select the plausible combinations of (*S*, *F*_msy_, *b*_ratio_) conditioned on the information from the stock assessments. Sixteen combinations of (*S*, *F*_msy_, *b*_ratio_) are retained by this operation (Table 1). We call these 16 combinations with estimated plausibility values ‘Bio-parameter Scenarios’. Reflecting a highly negative correlation between *S* and *F*_msy_ in Japanese fisheries stocks (Fig. 2), scenarios with high (low) *S* and high (low) *F*_msy_ are given low plausibility (Table 1). Bio-parameter Scenarios with high *b*_ratio_ are also given low plausibility reflecting a low frequency in Japanese stock assessment results. The Bio-parameter Scenarios with a combination of low *S*, low *F*_msy_ and high *b*_ratio_ are excluded because of extremely low plausibility (Table 1 and Fig. 2). Given *S*, *F*_msy_ and *b*_ratio_, the growth parameter necessary for the delay-difference model is estimated by solving equations under the equilibrium assumption (Appendix S3).

**Figure 2.**
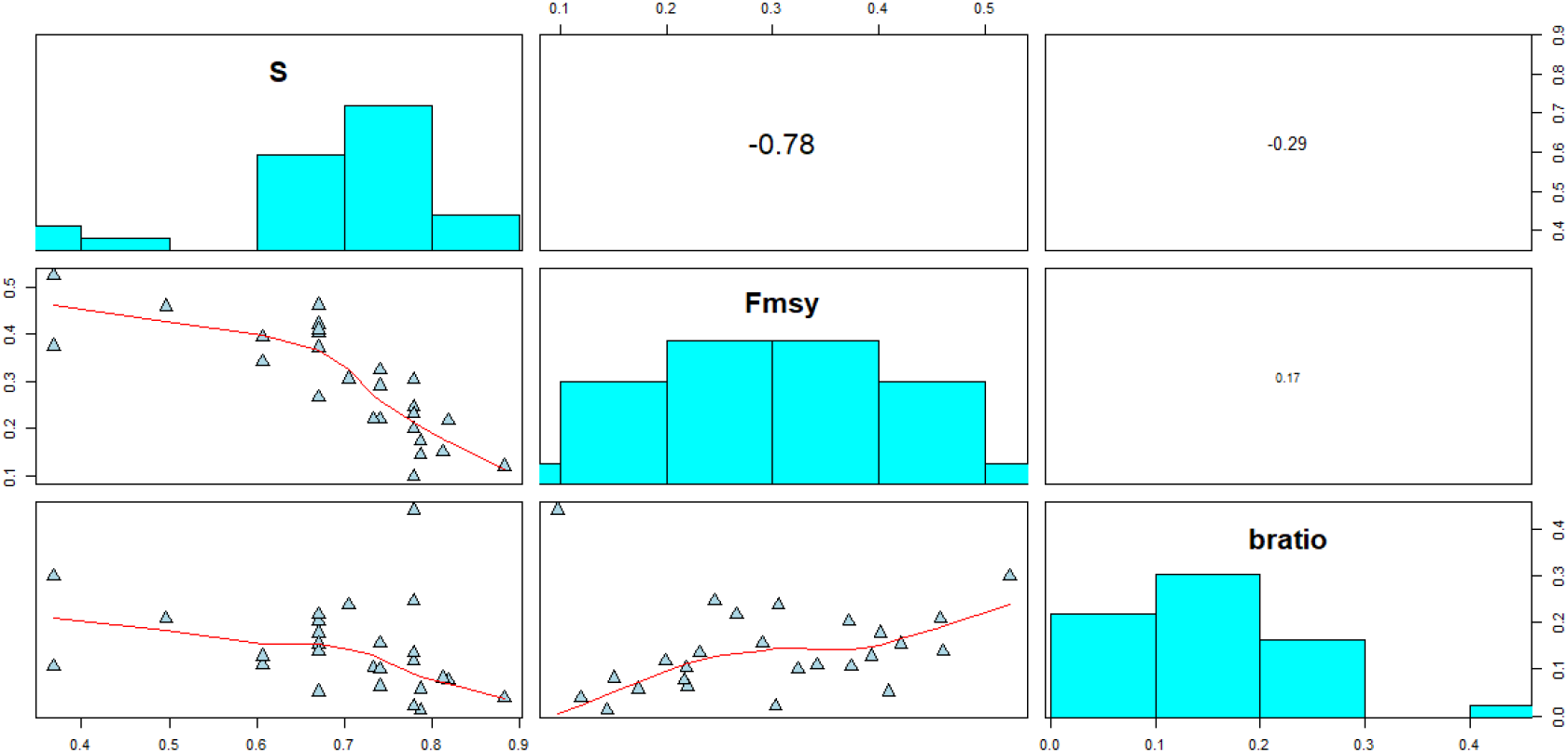
The biological parameters (natural survival rate (*S*), *F*_msy_ and *b*_ratio_) for 26 Japanese data-rich stocks. *F*_msy_ and *b*_ratio_ are estimated using a deterministic population dynamics model without assuming variation in R.

**Table 1.** Bio-parameter Scenarios.

| Bio-parameter |  |  |  |  |
| --- | --- | --- | --- | --- |
| Scenario | Survival Rate | $F_{msy}$ | $b_{ratio}$ | Plausibility |
| 1 | 0.82 | 0.10 | 0.10 | 0.075 |
| 2 | 0.67 | 0.10 | 0.10 | 0.004 |
| 3 | 0.82 | 0.25 | 0.10 | 0.198 |
| 4 | 0.67 | 0.25 | 0.10 | 0.208 |
| 5 | 0.82 | 0.40 | 0.10 | 0.007 |
| 6 | 0.67 | 0.40 | 0.10 | 0.168 |
| 7 | 0.82 | 0.10 | 0.25 | 0.032 |
| 8 | 0.67 | 0.10 | 0.25 | 0.004 |
| 9 | 0.82 | 0.25 | 0.25 | 0.059 |
| 10 | 0.67 | 0.25 | 0.25 | 0.148 |
| 11 | 0.82 | 0.40 | 0.25 | 0.001 |
| 12 | 0.67 | 0.40 | 0.25 | 0.083 |
| 13 | 0.37 | 0.40 | 0.25 | 0.003 |
| 14 | 0.82 | 0.25 | 0.40 | 0.001 |
| 15 | 0.67 | 0.25 | 0.40 | 0.007 |
| 16 | 0.67 | 0.40 | 0.40 | 0.003 |

In the reference case, we set the coefficient of variation in the deviance of log R (CV_R_) to 0.4 following Ichinokawa et al. (2017) and assume no observation error in biomass and no autocorrelation in R. The simulation is run for 50 years with a constant harvest rate before introducing HCR management. The constant harvest rate is determined by altering the level of SB/SB_msy_ after 50 years. SB (=B(1−F)) is projected from SB_0_ to SB_msy_ on average for the first 10 years and this projection (the first 10 years) is eliminated when estimating the SR relationship in the HCR. SB is then projected from SB_msy_ to 1) 0.2×SB_msy_ (Trajectory Scenario L), 2) 1.0×SB_msy_ (Trajectory Scenario M) and 3) 1.8×SB_msy_ (Trajectory Scenario H) on average after 40 years. Thus, we have 40 years of SR data at the start of management in which SB changed from SB_msy_ to 0.2, 1.0, or 1.8 times SB_msy_ on average. After the introduction of management, ABC calculated by HCR is subtracted from the stock biomass every year. The HCR-based management is then continued over 50 years. When calculating ABC, biological parameters other than the parameters of the HS curve are assumed to be known. The parameters of the HS curve are estimated every five years after the introduction of HCR management and necessary BRPs are calculated by simulation. All simulations for each scenario (i.e. a combination of Bio-parameter and Trajectory Scenarios) are iterated 100 times to estimate stochastic BRPs, stock biomass, and ABC.

#### 2–3–2. Sensitivity test trials

The uncertainty considered in the reference case is only the variation in R. We take other uncertainties into account as sensitivity test trials (Table 2). When the reference case (CV_R_ = 0.4) is the first scenario (Sensitivity Scenario 1), the second scenario (Sensitivity Scenario 2) enlarges the variation in R to 0.75 (CV_R_ = 0.75), which is almost the same as (actually slightly larger than) the global average (Thorson et al. 2014). The third and fourth sensitivity test (Sensitivity Scenarios 3 and 4) add the estimation error of biomass and SB using a log normal distribution LN(0, CV_obs_) (CV_obs_ = 0.2 for Sensitivity Scenarios 3 and CV_obs_ = 0.4 for Sensitivity Scenario 4) in calculating ABC and estimating the parameters of the HS curve. The fifth sensitivity test (Sensitivity Scenario 5) adds autocorrelation in R (*ρ* = 0.5), in which the value of autocorrelation is slightly larger than those of Thorson et al. (2014) (*ρ* = 0.43) and Ichinokawa et al. (2017) (*ρ* = 0.36). Although the second to fifth scenarios change only one factor at a time from the reference scenario to examine an impact of each factor, the sixth and seventh sensitivity tests (Sensitivity Scenarios 6 and 7) are combinations of several sensitivity tests. Sensitivity Scenario 6 has relatively moderate uncertainties with CV_R_ = 0.4, CV_obs_ = 0.2, and *ρ* = 0.5, whereas Sensitivity Scenario 7 has the worst uncertainties with CV_R_ = 0.75, CV_obs_ = 0.4, and *ρ* = 0.5.

**Table 2.** Sensitivity Scenarios.

| Sensitivity Scenario | Content | Source |
| --- | --- | --- |
| 1 | Reference set: $CV_R=0.4$, $CV_{obs}=\rho=0$ | Ichinokawa et al. (2017) |
| 2 | Large CV in recruitment: $CV_R=0.75$, $CV_{obs}=\rho=0$ | Thorson et al. (2014) |
| 3 | Small estimation CV in biomass: $CV_R=0.4$, $CV_{obs}=0.2$, $\rho=0$ | Okamura et al. (2017), Nishijima et al. (2019) |
| 4 | Large estimation CV in biomass: $CV_R=0.4$, $CV_{obs}=0.4$, $\rho=0$ | Nishijima et al. (2019), Yukami et al. (2019) |
| 5 | Autocorrelation in recruitment: $CV_R=0.4$, $CV_{obs}=0$, $\rho=0.5$ | Ichinokawa et al. (2017), Thorson et al. (2014) |
| 6 | All factors (moderate scenario): $CV_R=0.4$, $CV_{obs}=0.2$, $\rho=0.5$ | - |
| 7 | All factors (worst-case scenario): $CV_R=0.75$, $CV_{obs}=0.4$, $\rho=0.5$ | - |

#### 2–3–3. Performance measures

We calculate the following five quantities to investigate a performance of each HCR and compare relative performances among the three HCRs (Table 3). The performance measures for each HCR are standardized by dividing them by the performance measures for the constant 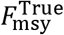 harvest control (i.e., 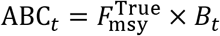 regardless of the stock status, and 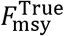 is calculated by the stochastic simulation from the true SR curve without any curve fitting process).

1. The average SB for the last 10 years divided by SB_msy_ (SB)
2. The average ABC for the last 10 years divided by MSY (CAT.L)
3. The average annual variation in ABC for last 10 years (AAV)
4. The average ABC for the first 5 years divided by MSY just after the introduction of management (CAT.F)
5. The average ABC for the first 5 years just after the introduction of management divided by the average catch for the last 5 years before the introduction of management (CC.F)

**Table 3.**
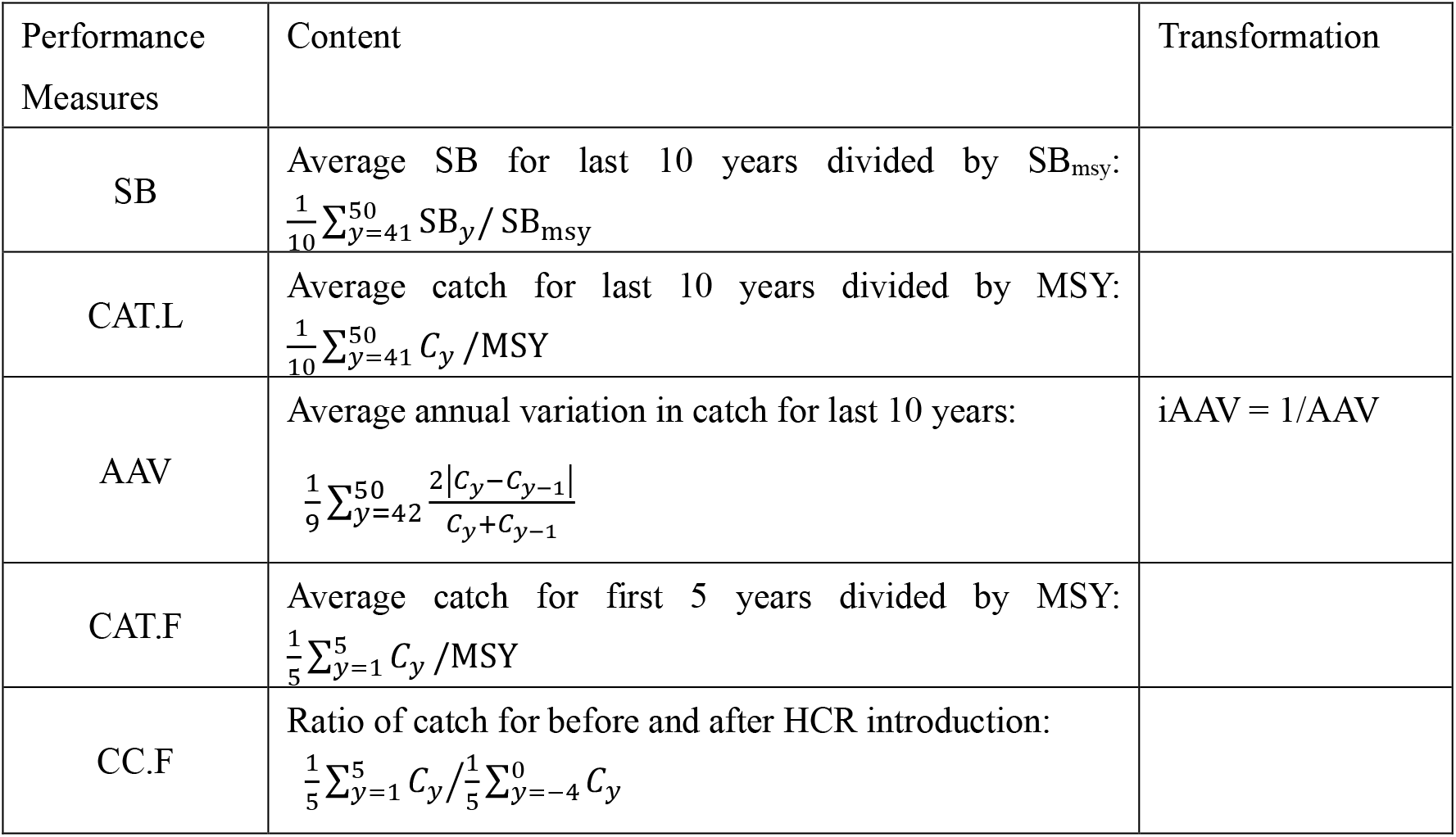
Performance Measures.

The first three performance measures are evaluations of the last stage of management (41–50 years after the introduction of HCR management), and they assess the status of population size, the amount of catch, and the variation in catch. The last two performance measures are evaluations of the early stage of management (1–5 years after the introduction of HCR management), and they assess the amount of catch, and the decreasing rate of catch before and after the management introduction. AAV is calculated by

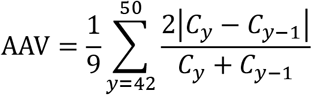

where *y* is the number of years after the introduction of management. When sequential catches in AAV become zero, they are simply eliminated as missing data. The performance measures are averaged by weighting over the plausibility of the Bio-parameter Scenarios.

The performance measures other than AAV are based on ‘bigger is better’, but AAV is based on ‘smaller is better’. We therefore transform AAV to ‘bigger is better’ by transforming to the inverse of AAV (iAAV = 1/AAV; Table 3).

We summarize the impacts of the factors (HCRs, CV_R_, CV_obs_, *ρ*, presence/absence of time lag, and whether the SR relationship is linear at the introduction of management) on each performance measure using the multiple linear regression models. The last factor is to determine which performance measures are affected when density dependence is not clearly detected over the observed stock sizes. We determine that the SR relationship is linear if the break point of HS is greater than 0.99 times the maximum value of historical SBs.

We then investigate how long each HCR takes to recover SB_msy_ for the Trajectory Scenario L (SB_msy_ → 0.2SB_msy_). The results are summarized by the annual probabilities that SB exceeds SB_msy_ for 50 years after the introduction of HCR management. We also investigate how the recovery time to SB_msy_ changes by the change of *β* for the new HCR.

For further confirming the robustness of HCR, additional sensitivity simulation tests are conducted for only the Trajectory Scenario L: i) the autocorrelation in the biomass estimates (*ρ*_obs_=0.5); ii) the random errors in ABC (CV_ABC_=0.2); iii) the systematic bias in the biomass estimates (B_obs_=1.5B_true_, SB_obs_=1.5SB_true_); iv) the true SR curve different from HS (the Beverton-Holt SR curve); and v) the true SR curve different from HS (the Ricker SR curve) (Appendix S4).

### 2–4. Forecast and hindcast analysis

Although the above analyses are based on a ‘typical’ stock’s parameters by conditioning on the real data of 26 stocks used in Ichinokawa et al. (2017), we also use real data more directly to demonstrate the effect and validity of the new HCR in terms of forecast and hindcast simulations. For forecasting, we use a delay-difference model similar to the MSE simulation and generate the historic data for each stock using the actual residuals of Rs. In future projections, the new Rs are generated from the bias-corrected HS SR function with the coefficients of variation for log Rs estimated from the stock assessment data. For hindcasting, we also use a delay-difference model similar to the MSE simulation, assuming the new HCR was introduced 5 or 10 years ago. We use 5 years when the length of the SR time series is less than 20 years, otherwise we use 10 years. The performance of NEW and NEW2 is compared with that of the actual SB and fishing rate. The details of the forecast and hindcast analyses are given in Appendix S5.

## 3. Results

We mention the results of the Trajectory Scenario L for the basic MSE simulation because the results in each HCR of this scenario show the greatest contrasts and are the most interesting. The graphical and tabular results of the Trajectory Scenarios M and H are given in Appendix S6 and briefly mentioned in the text.

A radar chart with or without a time lag displays the relative values of the five performance measures under Trajectory Scenario L (Fig. 3, Appendix S6). The circles with a radius of 1 are the results of the reference HCR that uses 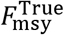 without a time lag. Because the results for some scenarios are similar, we focus on the Sensitivity Scenarios 1 (basecase), 4 (the largest observation error), and 7 (the worst case) (Fig. 3). The results for all Sensitivity and Trajectory Scenarios are given in Appendix S6.

**Figure 3.**
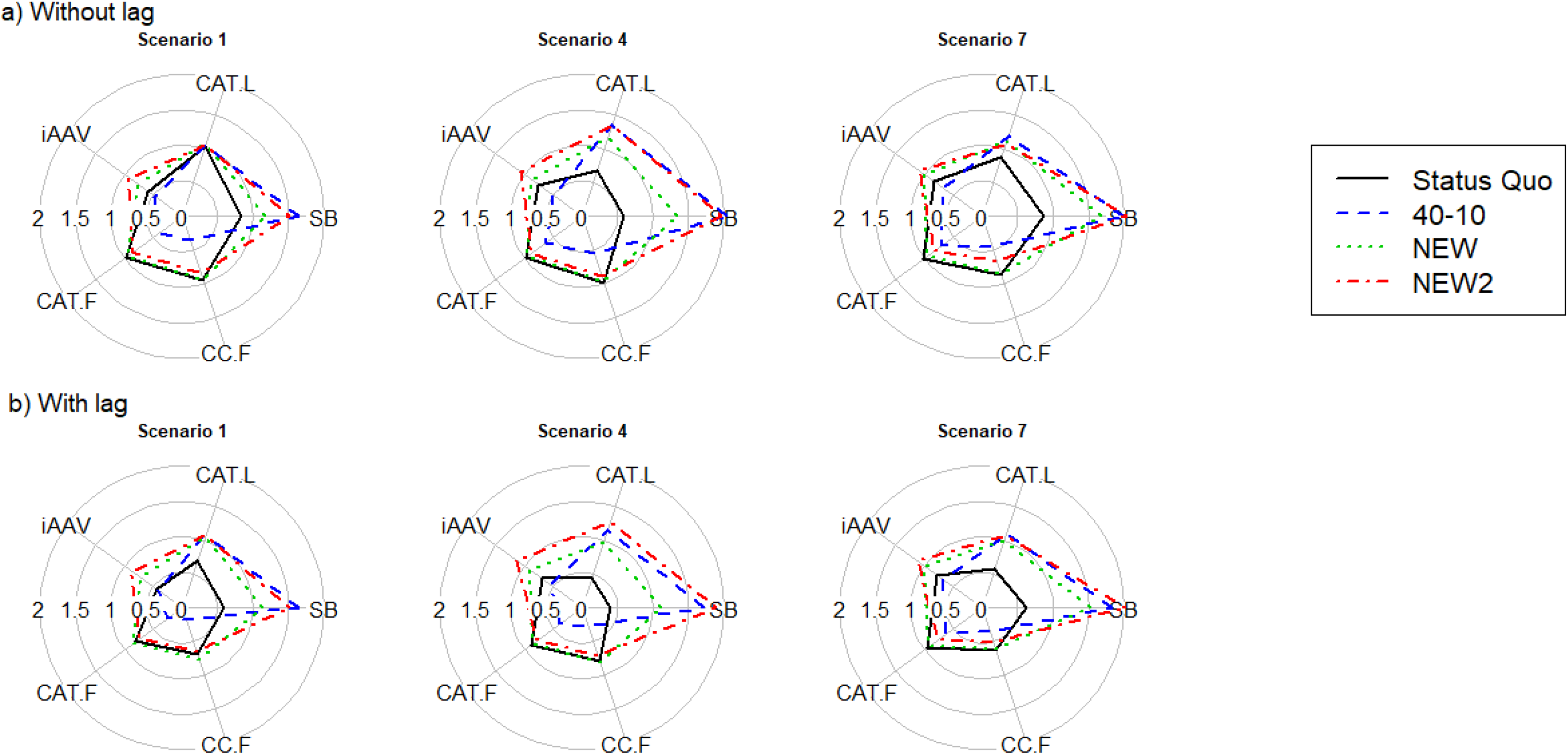
The radar chart plots of performance measures for Trajectory Scenario L over Sensitivity Scenarios; a) no time lag and b) a time lag. The explanation of performance measures is provided in Table 3. Status Quo is the traditional HCR in Japan, 40–10 is the 40–10 rule, NEW is the new HCR with *β*=1.0, and NEW2 is the new HCR with *β*=0.8. The circle with radius 1 denotes the results harvested by the true *F*_msy_.

When there is no time lag (Fig. 3a and Appendix S6), Status Quo shows the best performance measures in the early stage of management (CAT.F and CC.F). However, Status Quo often has the worst performance measures on SB and for the amount of ABC in the last stage of management (SB and CAT.L). Whereas 40–10 generally performs the best in SB, it has the worst in CAT.F, CC.F, and iAAV. NEW achieves the best balance among all performance measures, but the performance on SB are inferior to that of 40–10. NEW2 shows similar performance to NEW, but the performance on SB are comparable to that of 40–10, with a small sacrifice of performance in CAT.F and CC.F (Fig. 3a).

When there is a time lag of one year (Fig. 3b and Appendix S6), the polygons of Status Quo obviously become smaller compared to those without time lag. On the other hand, 40–10, NEW, and NEW2 are insensitive to whether there is a time lag. 40–10 generally shows the highest conservation performance with a sacrifice of performance in CAT.F and CC.F, and NEW and NEW2 have much larger iAAVs (or smaller AAVs) than the other two HCRs and are generally well-balanced over the five performance measures in all Sensitivity Scenarios (Fig. 3b). The conservation-related performance (SB) of NEW2 is almost equal to that of 40–10.

The results of Trajectory Scenario M and Trajectory Scenario H are more insensitive to the presence/absence of time lag than is Trajectory Scenario L (Appendix S6). In Trajectory Scenario M, the polygons are mostly round. The conspicuous differences among HCRs are detected in SB and AAV. NEW2 is the best for AAV, and 40–10 is the best for SB, while the conservation performance of 40–10 and NEW2 are quite similar (Appendix S6). Trajectory Scenario H provides shapes of polygons pointed to SB. In contrast to Trajectory Scenario L, Status Quo is the best performer on conservation but the worst performer on measures in the early stage of management. NEW2 is the best performer on AAV. NEW and NEW2 are well-balanced over all Sensitivity Scenarios (Appendix S6).

The impacts of factors on the five performance measures are quantified by multiple regressions for the Trajectory Scenario L (Fig. 4). Status Quo achieves considerably low SB and loses substantial CAT.L, while it shows the best performance for CAT.F. 40–10 achieves SB higher than SB_msy_ and almost MSY, at the sacrifice of high AAV and low CAT.F. NEW almost achieves SB_msy_ and MSY. NEW2 has performances similar to 40–10 for SB and CAT.L, while it has much smaller AAV than 40–10 and the smallest AAV among HCRs. NEW has almost the same CAT.F as Status Quo, whereas NEW2 has slightly lower CAT.F. The effect of CV_obs_ has the worst impact on performance measures in the last stage of management, indicating the importance of improving the precision of population biomass estimates. The presence of time lag generally has mild impacts. Whether the SR relationship is linear has weak effects on SB, CAT.L, and AAV, but a relatively large effect on CAT.F. This is probably because the linear SR relationship can easily take place when *b*_ratio_ is high (this indicates that more conservative management is needed).

**Figure 4.**
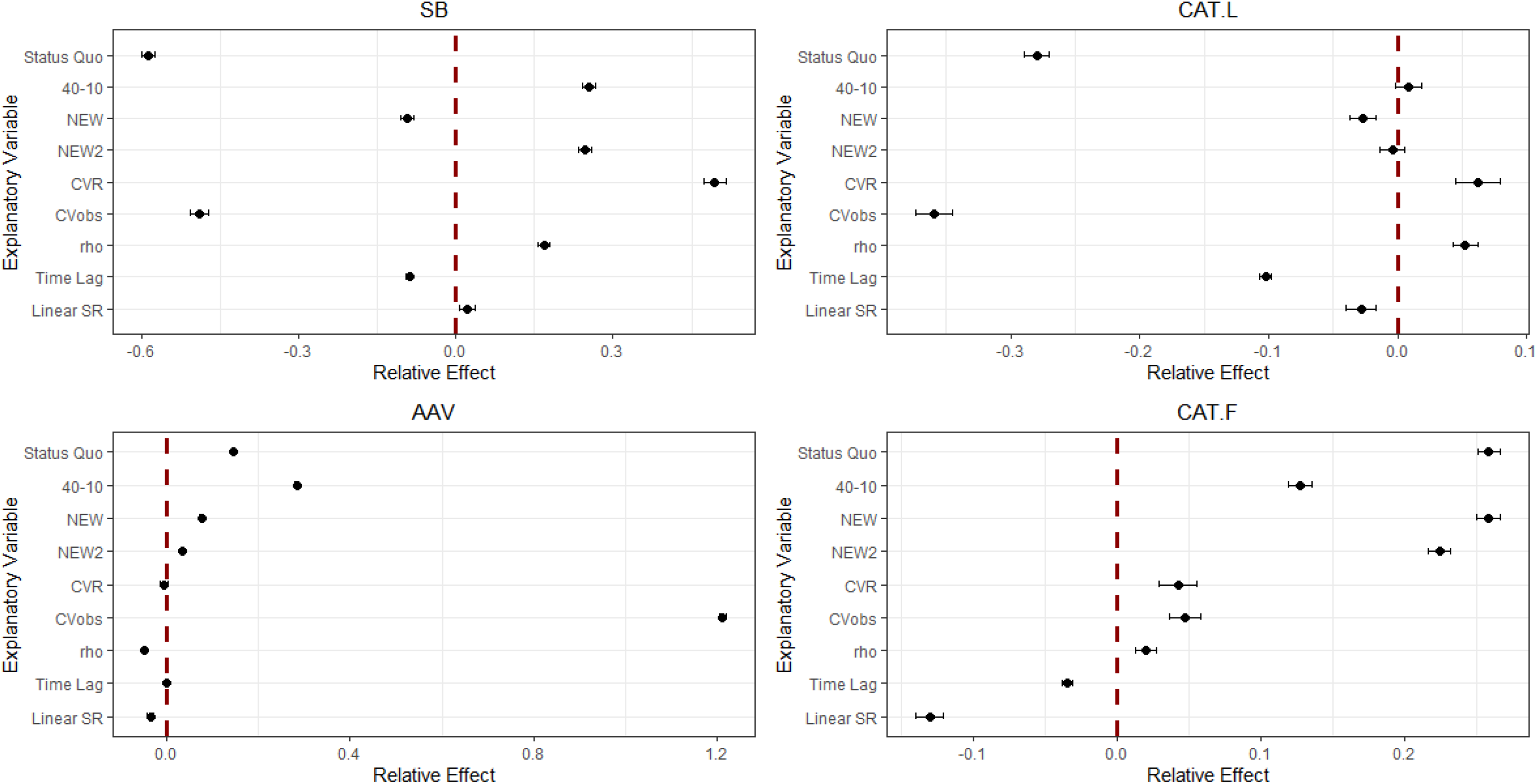
The relative effect of HCRs and uncertainty factors on performance measures. For SB and CAT.L, the effect of HCRs is evaluated by (SB_L_ − SB_msy_)/SB_msy_ and (C_L_ − MSY)/MSY where ‘L’ denotes the average for final 10 years in the simulation. Status Quo is the traditional HCR in Japan, 40–10 is the 40–10 rule, NEW is the new HCR with *β*=1.0, and NEW2 is the new HCR with *β*=0.8. CVR (=0.4, 0.75) is the CV of R, CVobs (=0, 0.2, 0.4) is the CV of biomass, *ρ* (=0, 0.5) is the autocorrelation of R, Time Lag is the presence/absence of time lag, and Linear SR is whether the SR relationship is linear at the introduction of management.

Status Quo is unsuccessful in the recovery to SB_msy_ in Trajectory Scenario L with a time lag (Fig. 5). NEW is unable to achieve the recovery to SB_msy_ in Sensitivity Scenario 4 (CV_obs_=0.4), and achievement in the other Sensitivity Scenarios takes 30 to 50 years, except for Sensitivity Scenario 2 (CV_R_=0.75), which takes just 20 years. The 40–10 achieves the recovery to SB_msy_ in 10 years except for in Sensitivity Scenario 7, and it takes 20 years until recovery for Sensitivity Scenario 7, which is the worst-case scenario including all large uncertainties. NEW2 makes slightly slower progress in comparison to 40–10, but generally it achieves the recovery to SB_msy_ in 10 years in most Sensitivity Scenarios, although it has a slightly lower than 50% probability. NEW2 also takes 20 years until recovery in Sensitivity Scenario 7. Sensitivity tests on *β* show that the population trajectories over all scenarios except Sensitivity Scenario 7 exceed SB_msy_ with a probability of about 0.5 in 10 years if *β* is equal to and less than 0.8 (Appendix S7).

**Figure 5.**
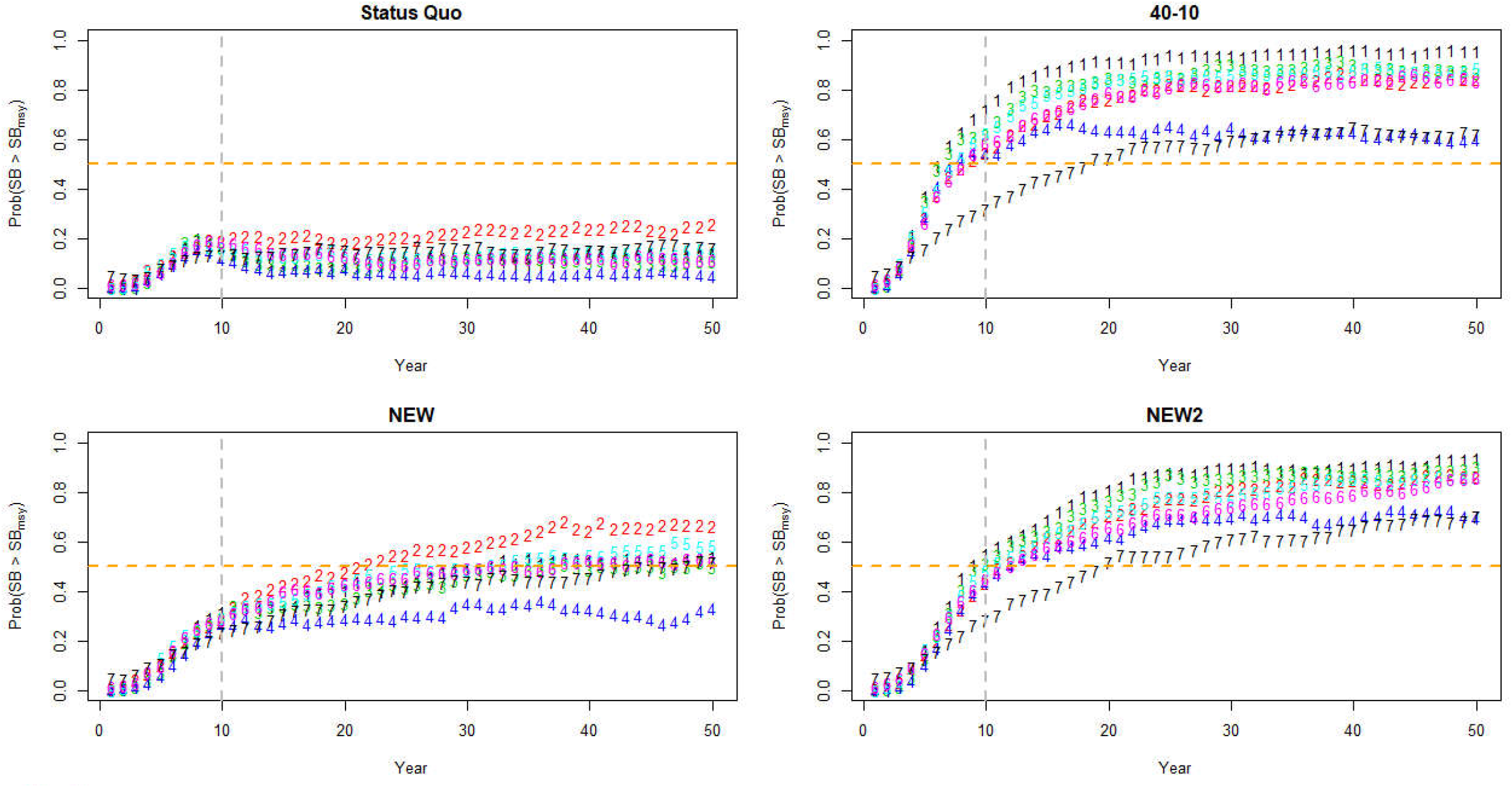
The trajectories of SB over SB_msy_ for Sensitivity Scenarios when there is a time lag. The horizontal broken line denotes the line at which Prob(SB > SB_msy_)=0.5 and the horizontal broken line denote 10 years after the start of HCR management.

The results for the Additional Sensitivity Scenarios are similar to those of the Sensitivity Scenarios in the sense that NEW/NEW2 are well-balanced over all the performance measures (Fig. 6 and Appendix S8). For the scenario with the Beverton-Holt curve as the true SR curve, even 40–10 and NEW2 fail to achieve SB_msy_ on average. However, the losses in the final catches from MSY are very low for both HCRs, reflecting the PGY concept (Hilborn 2010). Although 40–10 generally shows the highest conservation performance, substantial catch reduction needs to be accepted at the introduction of management.

**Figure 6.**
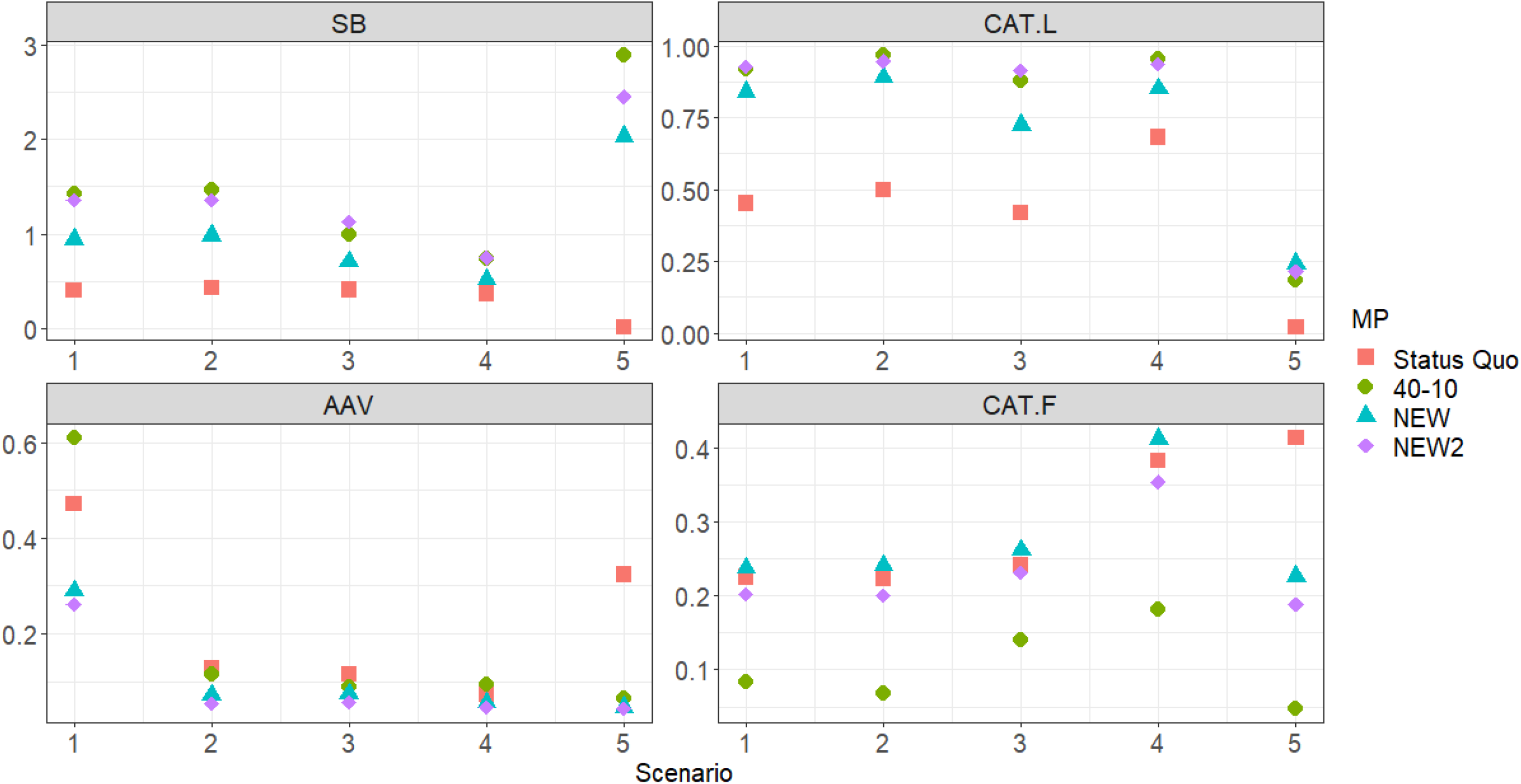
Comparison of main performance measures for Additional Sensitivity Scenarios when there is a time lag. Scenario 1: ρ_obs_ = 0.5, Scenario 2: CV_ABC_ = 0.2, Scenario 3: B_obs_ = 1.5B_true_ & SB_obs_ = 1.5SB_true_, Scenario 4: SR = BH, and Scenario 5: SR = RI.

In future projections, NEW2 shows a similar performance to 40–10 and has the lowest probability of overfishing (*F* > *F*_msy_) and overfished (SB < SB_msy_) among all HCRs (Fig. 7). Although NEW generally reduces overexploitation, the recovery of the stock status is insufficient. Status Quo fails to prevent overfishing and to rebuild overfished populations in most stocks even after 50 years. Although NEW2 mildly reduces fishing pressure compared to Status Quo at the onset, 40–10 drastically reduces fishing pressure, whereas the final stock statuses of NEW2 and 40–10 are similar (Appendix S9). For hindcasting, NEW2 provides substantially reduced fishing pressure compared to the actual fishing rate. Consequently, it reduces the probability of overfishing and overfished populations. However, the recovery of stock status is insufficient because of time shortages indicating that longer times are needed for the recovery of most stocks (Fig. 8).

**Figure 7.**
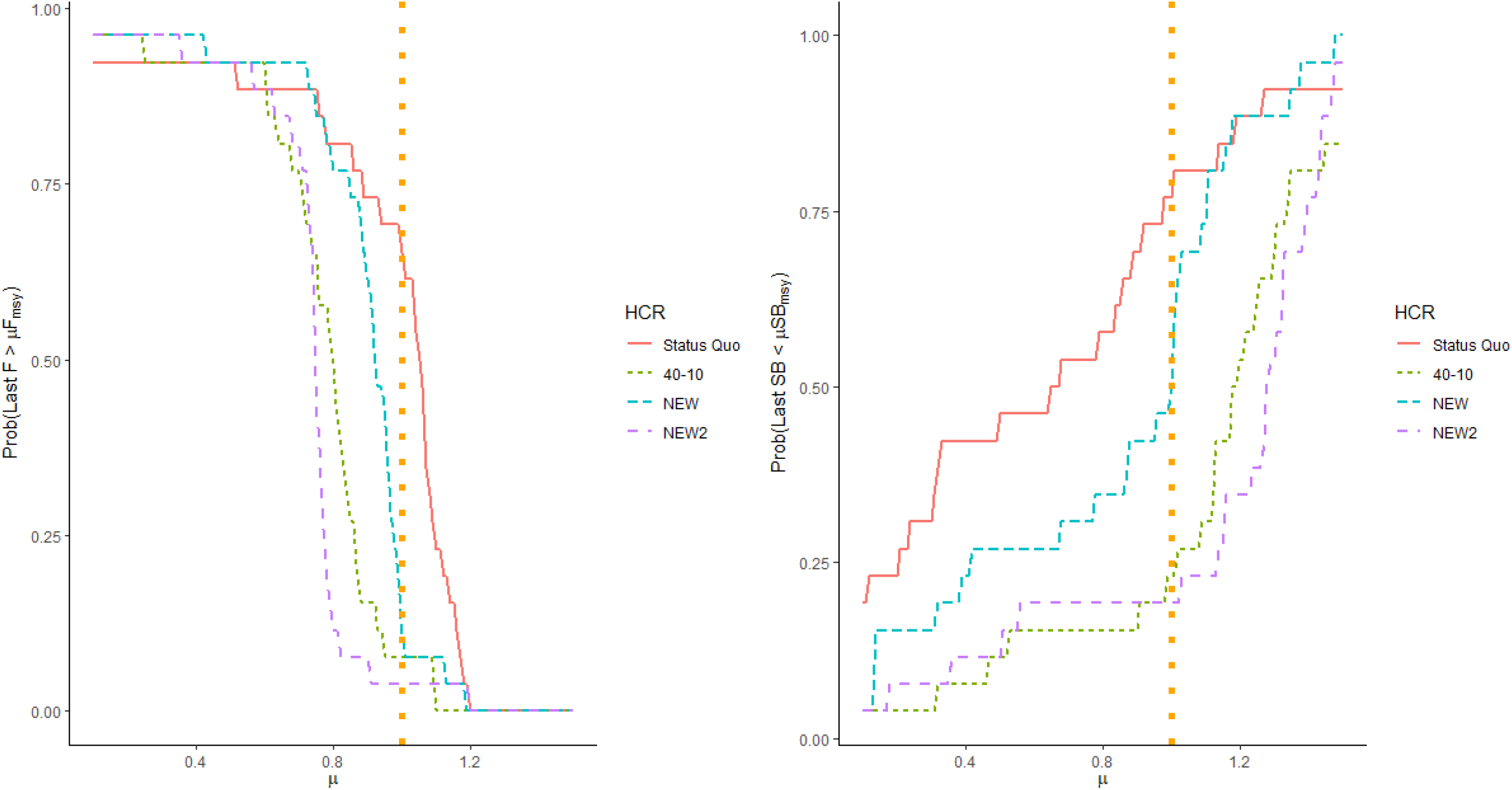
The probabilities that the last fishing rate is greater than μ×F_msy_ and the last SB is lower than μ×SB_msy_ for future projections (0 < μ < 1). The probabilities are calculated by the proportion that meets the condition out of 26 stocks.

**Figure 8.**
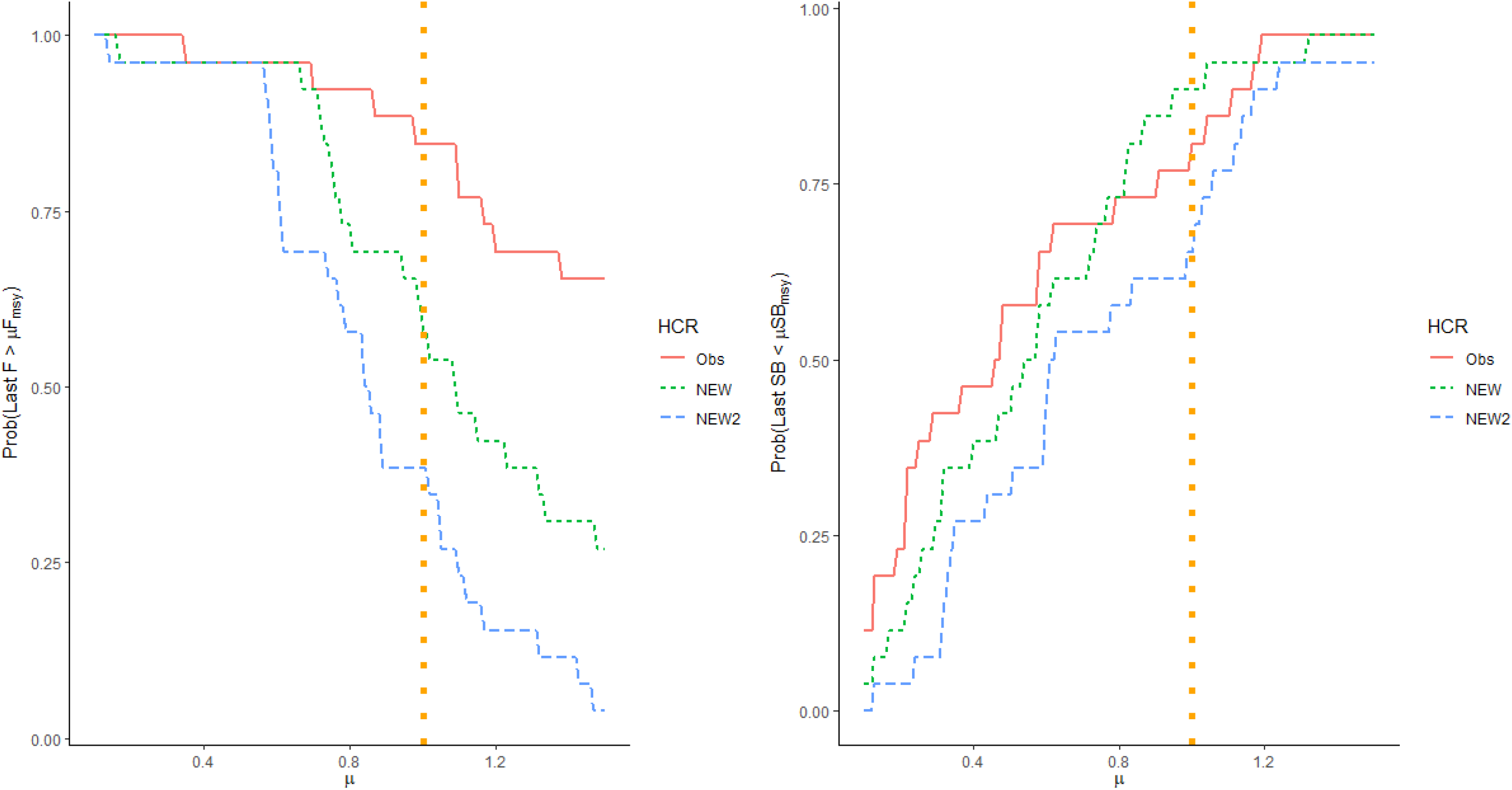
The probabilities that the last fishing rate is greater than μ×*F*_msy_ and the last SB is lower than μ×SB_msy_ for hindcastings (0 < μ < 1). The probabilities are calculated by the proportion that meets the condition out of 26 stocks.

## 4. Discussion

If we used the Beverton-Holt or the Ricker model as a SR curve in the new HCR, SB_msy_ and *F*_msy_ would sometimes become extremely low or high especially when the SR data are scarce or of low contrast causing low CAT.F. HS is able to circumvent such a problem using its characteristics (2). Furthermore, the stochastic SB_msy_ and *F*_msy_ from HS match the precautionary principle (the characteristics (3)) so that the management robust to large uncertainties is realized leading to stable catch (CAT.L close to MSY and low AAV) strengthened by high biomass (high SB). Because the R variation comes from various uncertainties including environmental changes, species interactions, and species’ biological characteristics, the inclusion into the HCR leads to benefits on ecosystem-based management in addition to achieving single species management objectives. The fact that NEW2 loses little sustainable yield but it has a higher conservation performance (Figs. 3 and 4) is consistent with the PGY concept (the characteristics (4)), indicating that NEW2 will have better economic and ecosystem outcomes.

Because maintaining a higher SB usually entails a lower fishing effort and hence, a lower cost, NEW2 is recommended as the best HCR from the standpoint of sustainable use, conservation, stable fisheries, economic efficiency, fisherman compliance, and ecosystem integrity. However, the reduction of catch tends to be met with strong resistance by fishermen. NEW then becomes a candidate for HCR, if its slower recovery and slightly inefficient performance can be accepted by all stakeholders.

Although the time lag is detrimental to all performance measures, the effects do not seem particularly serious except in Status Quo. When the SR relationship is linear and does not show density dependence at the start of HCR management, the performance in the last stage of management does not deteriorate except for a slight decrease in catch amount, whereas the performance in the early stage of management becomes poorer. This is probably because the linear SR relationship takes place when the *b*_ratio_ is high. More reduction of the catch in the initial stage of management is justified when the SR relationship is linear, since the population with a high *b*_ratio_ is more vulnerable to fishing in general. A vague SR relationship does not suggest that management with explicit targets is unnecessary. Rather, these results demonstrate that management using target biomass is still meaningful and useful even under large environmental uncertainties (Szuwalski et al. 2015).

Status Quo based on the deterministic model (the ‘sb’ method for SB_lim_) and the empirical model (*F*_med_) is vulnerable to low stock levels and stochasticity. Status Quo is also sensitive to the initial biomass states (Appendix S6). 40–10 allows catching by *F*_msy_ when SB exceeds 40%SB_0_, reduces the fishing rate when SB drops below 40% SB_0_, and prohibits fishing when SB falls below 10%SB_0_. However, because the fisheries stocks in Japan tend to have low SB_msy_, of generally less than 40%SB_0_ (Fig. 2), harvesting by *F*_msy_ does not lead to 40%SB_0_ in principle and as a result, the fishing rate continues to be reduced from *F*_msy_. This inconsistency inherent in 40–10 is reflected in its AAV, which is the highest among HCRs (Figs. 3, 4, and Appendix S6). Similarly using 10%SB_0_ for SB_ban_ may be too conservative for most fisheries stocks in Japan, considering the stock status of Japanese fisheries stocks is not healthy at the moment (Costello et al. 2016; Ichinokawa et al. 2017). On the other hand, NEW and NEW2 determine SB_lim_ by a loss of sustainable yield based on the past SR data, they can keep less variable catch that is close to MSY. Since the observed SR data are updated every five years, even if the target is not within the observed SR ranges at the onset, the target approaches the unbiased one as the data are accumulated.

The Magnuson-Stevens Fishery Conservation and Management Act specifies that overexploited stocks must be rebuilt within ten years (Patrick and Cope 2014; Benson et al. 2016). Many fisheries stocks in Japan that have been depleted by overexploitation could be rebuilt using the NEW2 HCR within 10 to 20 years, with a probability of slightly under 50% recovery to SB_msy_ except in the worst-case scenario (Sensitivity Scenario 7). Moreover, 40–10 can achieve recovery to SB_msy_ with a greater than 50% probability, while sacrificing the high probability of an initial complete loss of catch. If the Japanese government aims at recovering depleted stocks as quickly as possible similar to the US and avoiding complete fishing bans, we recommend using a new management procedure with *β* equal to and less than 0.8 (Appendix S7).

Our simulation is generic and is used to select management procedures that are suitable for the biological characteristics of typical Japanese fisheries stocks. It is recommended to conduct further case-specific simulations for more efficient management (Butterworth and Punt 1999). The following areas still need to be examined: sensitivities for the assumption of selectivity-at-age (Ichinokawa et al. 2014; Okamura et al. 2014); alternative models for the SR relationship (Ichinokawa et al. 2017); the effects of the population dynamics model with multiple stable states forced by significant environmental changes (King et al. 2015); and multispecies/ecosystem effects (Okamura et al. 2018). A bias in selectivity at age affects the SB_msy_ estimate but not the SB_0_ estimate. While the NEW/NEW2 HCRs are influenced by biases in *F*_msy_ and SB_msy_, the 40–10 HCR is influenced by a bias in *F*_msy_ alone. Therefore, unless biases in *F*_msy_ and SB_msy_ are offset, the NEW/NEW2 would suffer more than the 40–10 would. Similarly, the basic biological parameters are assumed to be known in the model fitting in our simulation. This has the potential to influence NEW/NEW2 more so than 40–10. However, HS would be relatively robust to the uncertainty compared to other differentiable nonlinear SR curves, such as the Beverton-Holt and Ricker curves, as SB_msy_ is usually within or near the observed ranges of SB (the characteristics (2) and (3)). Although we tried to evaluate the effects of using different SR curves (Fig. 6, Appendix S8), this trial should be further extended and investigated. The uncertainty in the SR relationship is generally large, but if we use a model selection procedure such as AIC, the probability that the correct SR relationship will be selected gradually becomes larger. Consequently, the SB_msy_ based on the true SR relationship will be finally reached, even though it may take a long time (Ichinokawa, unpublished data). Because the Ricker model tends to be less conservative (Ichinokawa et al. 2017), using the HS model may not lose conservation performance when the true SR relationship is the Ricker model. The best model selection or the model averaging using AIC may be effective for coping with the uncertainty in the SR relationship (Buckland et al. 1997). The effects of the regime shift are partly dealt with by the autocorrelation in the residuals of log R. However, if there is an extremely large and abrupt change in the population status, and the population truly has multiple equilibria, incorporating the autocorrelation may not completely resolve the problem. When a future regime shift is accurately and precisely predicted, it would be incorporated into the HCRs. However, because the estimation of regime shifts often suffers from considerable uncertainty, the inclusion of regime shifts in HCRs and BRPs is usually unsuccessful (Punt et al. 2013; King et al. 2015). Currently, multispecies effects are not explicitly incorporated into our HCR. In particular, multispecies fisheries could emerge as an immediate problem in fisheries management in Japan. The concept of ‘pretty good multispecies yield’ would then be a helpful tool for determining the *F*_msy_ range, taking into account multispecies effects and several objectives with trade-offs (Rindorf et al. 2017).

Fisheries management in Japan is definitely progressing. The large potential for improvement in fisheries management will affect the real outcomes of conservation efforts as well as economy and ecosystem integrity. The selection and adoption of new harvest control rules based on management strategy evaluation is one way to accelerate progress. As a matter of fact, the management procedure based on the new HCR has been applied and implemented to some stocks (http://www.fra.affrc.go.jp/shigen_hyoka/SCmeeting/2019-1/index.html). Box (1976) stated that ‘All models are wrong’. In this sense, the models and concepts relating to marine ecosystems such as MSY are certainly wrong. However, we believe that some traditional fisheries management tools such as MSY-related BRPs can remain useful in sustainable fisheries if they are correctly implemented (Hilborn and Ovando 2014; Hilborn et al. 2020). The new HCR using an HS SR curve with a stochastic simulation has affinities with the precautionary principle and ‘post-normal’ science (Funtowicz and Ravetz 1993; Dankel 2016). In sum, the new HCR can become a useful tool in Japan’s fisheries management under conditions of large uncertainty.

## Acknowledgements

We appreciate valuable comments from Prof. Doug Butterworth, University of Cape Town.

