## Supplements for "Evaluating a harvest control rule to improve the sustainability of Japanese fisheries"

### Supporting Information

#### Appendix S1. HS can circumvent extrapolating excessively beyond the historical SB time series

To investigate the robustness of the HS SR curve, we conduct a simple simulation using the stochastic Allen-Clark delay-difference model (Quinn and Deriso 1999)

$$N_{t+1} = S_t N_t + R_{t+1} e^{\varepsilon_t - 0.5\sigma^2}$$

where $S_t = \exp(-M - F_t)$, $M = 0.4$, $\varepsilon_t \sim N(0, \sigma^2)$, and $\sigma = 0.4$. $R_t$ is the Beverton-Holt (BH) SR relationship $R_t = aN_t(1 - F_t)/[1 + N_t(1 - F_t)/b]$, where we assume that $a = 1.2$ and $b = 1000$. We generate the dataset with 100 SB and R pairs and fit the HS SR curve and the BH SR curve to the generated data. For the HS SR curve, we set the maximum SB to the break point of HS when the SR curve is estimated to be linear, whereas we set the minimum SB to the break point of HS when the SR curve is estimated to be flat (Fig. S1–1). The fitted data are limited to the subsets (SB < 350, SB < 800, SB < 1200, and SB < 1500) of the full data with the maximum of SB = 1500 (Fig. S1–2). For the dataset limited to SB < 350, the BH curve goes to the very large value far outside the observed SB range, whereas the HS curve takes the break point to the maximum value of observed SBs. When the dataset is extended to SB < 800, the BH curve still takes a large value outside the observed SBs, though it comes close to the maximum value of SB. On the other hand, the HS curve has the break point within the observed SBs. As the dataset comes close to the full dataset, the BH curve becomes close to the truth while fluctuating, whereas (of course) the HS curve does not converge to the true curve, but shows strong robustness on the position of break point. This robustness of the break point of HS implies the robustness of $SB_{msy}$ estimated from the HS curve. The density-dependent parameter of BH fluctuates seriously depending on the sample size of the subsampled data, and easily tends to go far from the observed SB range when there is small sample size.

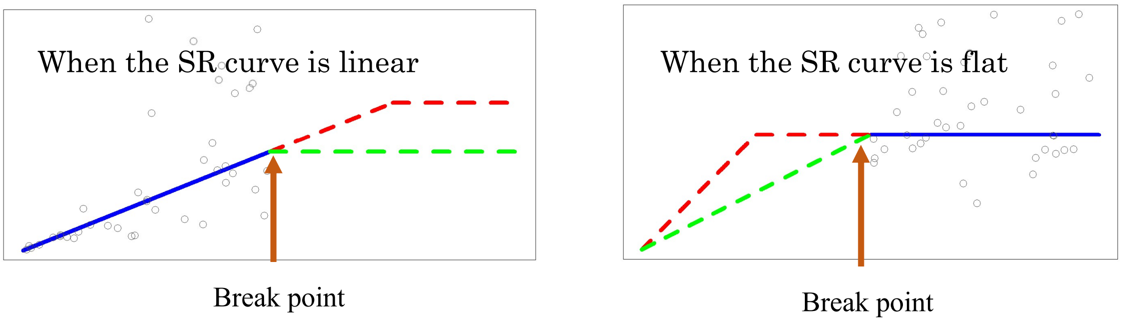


Fig. S1–1. The position of the break point of the HS curve when it is not statistically determined.

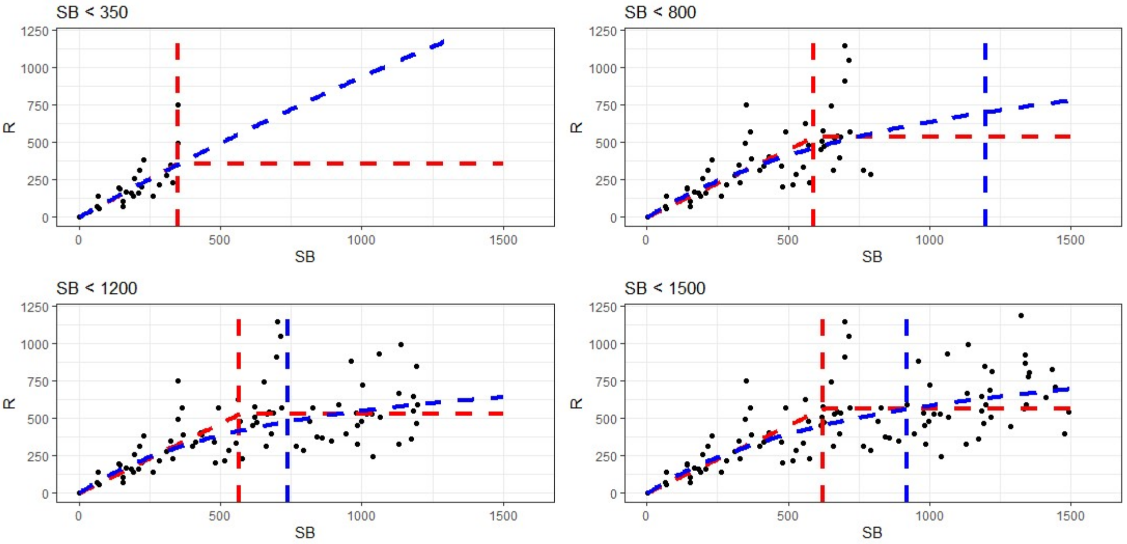


Fig. S1–2. The fitted SR curves (blue = BH, red = HS). The original dataset is generated with the sample size of 100 and the maximum value of SB is 1491.4. The SR curves are separately fitted to the subsampled datasets (SB < 350, SB < 800, SB < 1200, and SB < 1500 (full data)). The vertical lines denote the positions of the estimated $b$ parameters.

#### Appendix S2. Consistency of the HS SR curve with a precautionary approach

Here we investigate how the target BRP changes as the variation in R increases when the stochastic HS SR curve is used.

We use the stochastic Allen-Clark delay-difference model

$$N_{t+1} = S_t N_t + R_{t+1} e^{\varepsilon_t - 0.5\sigma^2}$$

where $\varepsilon_t \sim N(0, \sigma^2)$. $R_t$ is the HS curve

$$R_t = \begin{cases} aN_t(1 - F_t) & N_t(1 - F_t) \le b \\ ab & N_t(1 - F_t) > b \end{cases}$$

We assume that $S_t = \exp(-M - F_t)$, $M = 0.4$, $a = 1.2$, and $b = 1000$. Assuming constant $F$, we conduct the 500 years future projection with different $\sigma^2$ ( $\in \{0.0, 0.2, 0.4, 0.6, 0.8, 1.0\}$). Then we calculate the quasi-equilibrium biomass and catch after 500 years.

As the R variation σ increases, the equilibrium abundance and yield curve shift to left (Fig. S2–1) and as a result, the abundance at the MSY increases whereas the MSY and the fishing rate at MSY decrease. The analysis using the Pella-Tomlinson model showed the similar shift of MSY (Bordet and Rivest 2014). Because large recruitment uncertainty guarantees higher target abundance and lower fishing pressure (Figs. S2–1 and S2–2), the HS SR curve automatically enables managers to take the precautionary measures in advance.

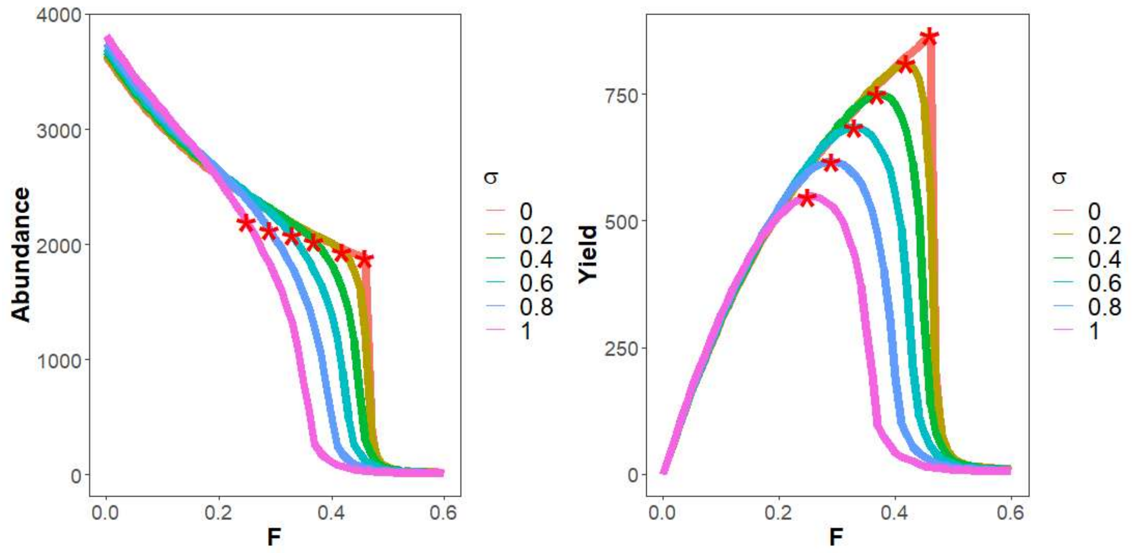


Fig. S2–1. Equilibrium abundance and yield for different σ under the HS SR curve. The star symbols correspond to the maximum sustainable yield.

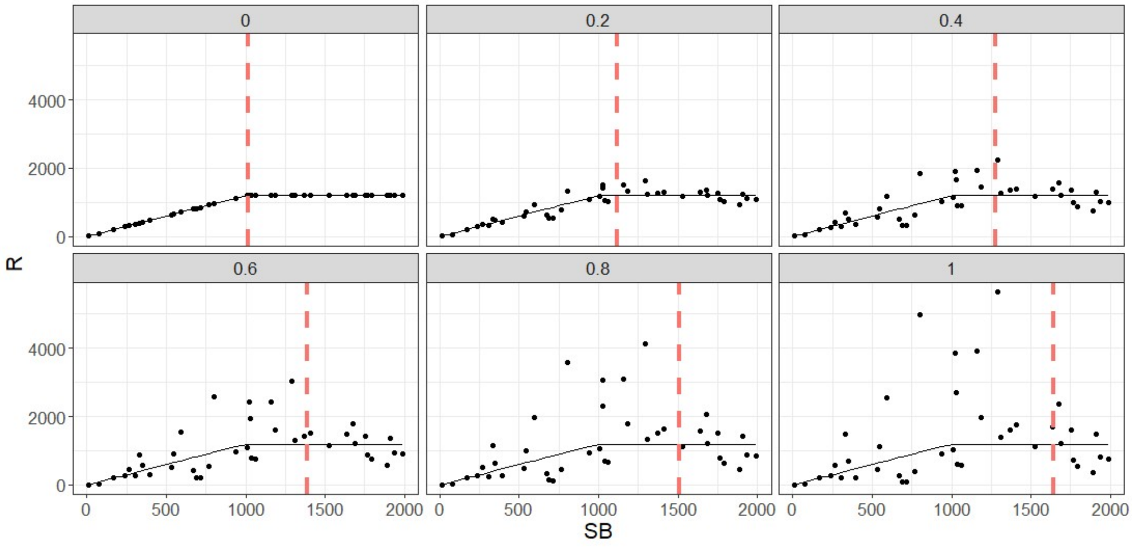


Fig. S2–2. The transition of SB for the change of the magnitude of R variation (σ = 0, 0.2, 0.4, 0.6, 0.8, and 1.0).

#### Appendix S3. The specification of population dynamic model used for generating the simulated data.

The population dynamics model is given by the general delay-difference model (Hilborn and Walters 1992; Quinn and Deriso 1999):

$$B_{t+1} = S(\theta N_t + \phi B_t)(1 - F_t) + w_r R_{t+1}$$

$$N_{t+1} = S N_t (1 - F_t) + R_{t+1}$$

where the recruitment $R_t$ is given by the stochastic HS curve:

$$R_t = \begin{cases} a\mathrm{SB}_{t-1} e^{\varepsilon_t} & \mathrm{SB}_{t-1} \le b \\ abe^{\varepsilon_t} & \mathrm{SB}_{t-1} > b \end{cases}$$

and $\mathrm{SB}_t = B_t(1 - F_t)$ is the spawning biomass. The residual of log recruitment $\varepsilon_t$ follows $\mathrm{N}\left(\rho\varepsilon_{t-1} - \frac{\sigma^2}{2(1-\rho^2)}, \sigma^2\right)$ where $\sigma^2$ is the variance and $\rho$ is the autocorrelation of log-recruitment (Thorson et al. 2015). The growth parameters $\theta$ and $\phi$ are derived from the linear relationship (a Ford-Walford plot):

$$w_a = \theta + \phi w_{a-1}$$

and $w_r$ is the mean weight at age at recruitment ($r$). We assume that $w_{r-1} = 0$ and $w_r = \theta$. The initial biomass $B_0$ without fishing is assumed to be equal to 10,000 without loss of generality.

Given deterministic $F_{\mathrm{msy}}$ and $b_{\mathrm{ratio}}$ (= $b/\mathrm{SB}_0$), we need to estimate the SR parameter $a$ and growth parameters $\theta$ and $\phi$. The break point $b$ is given by $\hat{b} = b_{\mathrm{ratio}}\mathrm{SB}_0 = b_{\mathrm{ratio}}B_0$ ($F_0$=0). Assuming the equilibrium under $F_{\mathrm{msy}}$ without stochastic error, the biomass at MSY is given by solving the above population dynamics model as follows:

$$B_{\mathrm{msy}} = \frac{\theta ab}{[1 - S(1 - F_{\mathrm{msy}})][1 - \phi S(1 - F_{\mathrm{msy}})]}$$

under the assumption of $w_r = \theta$. We cannot then differentiate between $\theta$ and $a$ such that we can assume $\theta = 1$ without loss of generality. Because MSY = $F_{\mathrm{msy}} \times B_{\mathrm{msy}}$, the Brody growth coefficient $\phi$ is determined by solving d MSY/d$F_{msy}$ = 0 given $F_{msy}$. Because this equation is a linear equation about $\phi$, the solution is provided by an analytical form:

$$\hat{\phi} = \frac{1 - S}{S[1 - S(1 - F_{msy}^2)]}$$

Note that the Brody growth coefficient $\phi$ is determined depending on only $S$ and $F_{msy}$ and unrelated to $\theta$, $a$, and $b$. Given $\hat{\phi}$, the parameter $a$ is derived from the equation of the biomass at equilibrium without fishing ($B_0$) as follows:

$$\hat{a} = B_0(1 - \hat{\phi}S)/\{[S/(1 - S) + 1]\theta b\}$$

The parameter $a$ depends on only $\hat{\phi}$, $S$, and $b_{ratio}$ because $B_0$ is cancelled out and now $\theta$ is fixed at 1. Given $\theta, \hat{a}, \hat{b}, \hat{\phi}$, $S$, and $F_{msy}$, $\mathrm{SB_{msy}} = B_{msy}(1 - F_{msy})$.

The ranges of basic biological parameters ($S$, $F_{msy}$ and $b_{ratio}$) are selected separately for each parameter from results using the HS curve in Ichinokawa et al. (2017): $S \in \{0.37, 0.67, 0.82\}$ (these values correspond to natural mortality coefficients $M$ = 1.0, 0.4, and 0.2, respectively), $F_{msy} \in \{0.10, 0.25, 0.40\}$, and $b_{ratio} \in \{0.10, 0.25, 0.40\}$. Considering correlations among biological parameters, we calculate a trivariate normal distribution N($\mu$, $\Sigma$) where $\mu$ is the mean vector of ($S$, $F_{msy}$, $b_{ratio}$) and $\Sigma$ is the corresponding variance-covariance matrix for 26 stocks in Ichinokawa et al. (2017). The probability densities are calculated by fitting the above combination of ($S$, $F_{msy}$, $b_{ratio}$) to the trivariate normal distribution. When the standardized probability density is less than 0.001, the combination is eliminated as it is too unrealistic. This operation produces 16 combinations with plausibility estimates in Table 1. All subsequent results in the MSE analyses are averaged by the plausibility weights.

The simulation is started from an equilibrium without stochasticity and fishing. Using the stochastic simulation with the variation in R, fishing is continued for 10 years with a constant fishing mortality adjusted such that SB after 10 years attains $\mathrm{SB_{msy}}$ on average.

This first 10 years data are eliminated from the analysis. The population is then projected from $SB_{msy}$ to $p \times SB_{msy}$ for 40 years with an adjusted constant fishing mortality rate where $p$ is set to 0.2, 1.0, and 1.8 (Trajectory Scenarios L, M, and H, respectively).

The 40-years data are used for fitting the HS curve to SR time series data with errors (i.e., the R variation as well as the SB variation) when the management by HCR is started. The BRPs ($SB_0$, $SB_{msy}$, $SB_{60\%msy}$, $SB_{10\%msy}$, and $F_{msy}$) are calculated by the stochastic simulation using the estimated SR parameters and the estimates of coefficient of variation (CV) and autocorrelation in R. The CV and autocorrelation in R are estimated using the maximum likelihood approach based on the exact likelihood (i.e. assuming the starting value is at the equilibrium state) by fitting the AR(1) model to the residuals of observed log R and predicted log R. The BRPs are updated every five years and ABC is calculated from HCR using the biomass information and the fishing rate every year. ABC is eliminated from the population dynamic model every year without implementation errors (However, see Additional Sensitivity Scenario 2 in Tables S4–1 of Appendix S4). If there is a time lag, the information on biomass in the latest year is excluded from the calculation of BRPs and ABC. When there is observation error in the Sensitivity Scenarios (Table 2), both of BRPs and ABC are calculated using the observed SB and biomass with errors. The simulation is conducted for 50 years after the start of HCR management. The performance measures (Table 3) are finally calculated and evaluated using the first 5years and the last 10 years of 50-year time series data on SB and ABC obtained from the simulation trials.

For Status Quo, ABC is calculated by multiplying $F$ by $B$ simply (without simulation). $SB_{lim}$ is updated every five years but $F_{med}$ is updated every year. For 40–10 and NEW/NEW2, ABC is calculated from the stochastic mean of $F \times B$ by one year forward simulation. $F_{msy}$, $SB_{lim}$, and $SB_{ban}$ are updated every five years.

#### Appendix S4. The specification of additional sensitivity scenarios.

We investigate the effects of autocorrelation in biomass estimates ($\rho_{\mathrm{obs}}$=0.5), variation in ABC or catch ($\mathrm{CV}_{\mathrm{ABC}}$=0.2 in log-normal distribution), overestimate in biomass ($\mathrm{B}_{\mathrm{obs}}$=1.5$\mathrm{B}_{\mathrm{true}}$, $\mathrm{SB}_{\mathrm{obs}}$=1.5$\mathrm{SB}_{\mathrm{true}}$) and true SR curves different from HS (the Beverton-Holt and Ricker SR curves) as Additional Sensitivity Scenarios (Table S4-1).

When the Beverton-Holt curve,

$$R_{t+1} = \frac{a\mathrm{SB}_t}{1 + \mathrm{SB}_t/b},$$

is used as the true SR curve, the $F_{\mathrm{msy}}$, $b_{\mathrm{ratio}}$, $\hat{\phi}$, and other biological parameters are assumed to be equal to those used in the simulations using the HS curve. Then, by solving the deterministic delay-difference equation with the Beverton-Holt SR curve, we obtain the slope parameter of the Beverton-Holt SR curve:

$$\hat{a} = \frac{(1 - \hat{\phi}S)(1 + b_{ratio}^{-1})}{w_r + \theta S/(1 - S)}.$$

Then the true population dynamics is projected using the stochastic simulation with the Beverton-Holt SR relationship. However, when estimating the SR parameters and the BRPs, we use the HS SR curve consistently for 40–10 and NEW/NEW2 HCRs.

We use the same procedure when using the Ricker curve.

Table S4–1. Additional Sensitivity Scenarios.

| Additional Sensitivity Scenario | Content |
|---|---|
| 1 | Autocorrelation in biomass: $CV_R$=0.4, $CV_{obs}$=0.2, ρ=0, $\rho_{obs}$=0.5 |
| 2 | CV in ABC: CVR=0.4, CVobs=0=ρ=0, $CV_{ABC}$=0.2 |
| 3 | Bias in biomass: $CV_R$=0.4, $CV_{obs}$=ρ=0, $B_{obs}=1.5\times B_{true}$ |
| 4 | Beverton-Holt stock-recruitment curve: $CV_R$=0.4, $CV_{obs}$=ρ=0 |
| 5 | Ricker stock-recruitment curve: $CV_R$=0.4, $CV_{obs}$=ρ=0 |

Appendix S5. The specification of future and hindcast prediction using the real stock assessment outputs.

The future projections using the real stock assessment outputs (26 stocks, Table S5–1) more directly are conducted using the delay-difference model:

$$B_{t+1} = S(\theta N_t + \phi B_t)(1 - F_t) + w_r R_{t+1}$$

$$N_{t+1} = S N_t (1 - F_t) + R_{t+1}$$

where the recruitment $R_t$ is given by the stochastic HS curve. We use the same assumptions as those used in the MSE simulation for the growth parameters and $B_0$, whereas we use the stock-specific survival rate, $F_{\text{msy}}$, and $b_{\text{ratio}}$. The deviation between the observed recruitment and the predicted recruitment is used to reproduce the trajectory of stock status in the past period for each stock. The initial SB is estimated using $B_t^{\text{obs}}/B_0^{obs} \times B_0$ which $B_t^{obs}$ is the biomass estimate obtained from the stock assessment and $B_0^{obs}$ is the initial equilibrium biomass estimate obtained from the stock assessment (In this case, virtual population analysis was used for all stock assessments).

For forecasting, we conduct the 50-years simulation with the HCR management under the estimated recruitment variation and autocorrelation. For hindcasting, we conduct the management with the new HCR for the last 5- and 10-years under the R variation and autocorrelation observed in the past (When the length of time series for SR relationship is less than 20 years, the 5 years back is adopted. Otherwise, the 10 years back is adopted).

Table S5–1. Twenty-six Japanese fishery stocks with SR data used for future and hindcast prediction.

| Stock ID | Common name | Scientific name | Region | Length of years |
|---|---|---|---|---|
| PILCHPJPN | Japanese sardine | *Sardinops melanostictus* | Pacific | 21 |
| PILCHTSST | Japanese sardine | *Sardinops melanostictus* | Tsushima Warm Current | 16 |
| JMACKPJPN | Japanese jack mackerel | *Trachurus japonicus* | Pacific | 32 |
| JMACKTSST | Japanese jack mackerel | *Trachurus japonicus* | Tsushima Warm Current | 41 |
| CMACKPJPN | Chub mackerel | *Scomber japonicus* | Pacific | 33 |
| CMACKTSST | Chub mackerel | *Scomber japonicus* | Tsushima Warm Current | 41 |
| SMACKPJPN | Spotted mackerel | *Scomber australasicus* | Pacific | 19 |
| SMACKECS | Spotted mackerel | *Scomber australasicus* | Tsushima Warm Current | 22 |
| APOLLPSOJ | Walleye pollock | *Gadus chalcogramma* | Sea of Japan | 34 |
| APOLLPJPN | Walleye pollock | *Gadus chalcogramma* | Pacific | 33 |
| RHERRTSST | Round herring | *Etrumeus teres* | Tsushima Warm Current | 38 |
| JANCHOPJPN | Japanese anchovy | *Engraulis japonicus* | Pacific | 36 |
| JANCHOSETO | Japanese anchovy | *Engraulis japonicus* | Seto Inland Sea | 37 |
| AMBERJ | Amberjack | *Seriola quinqueradiata* | Around Japan | 20 |
| RBRMSETOE | Red seabream | *Pagrus major* | Seto Inland Sea (east) | 37 |
| RBRMSETOW | Red seabream | *Pagrus major* | Seto Inland Sea (west) | 37 |
| RBRMECS | Red seabream | *Pagrus major* | Sea of Japan and East China Sea | 28 |
| SPANMACKSETO | Japanese Spanish mackerel | *Scomberomorus niphonius* | Seto Inland Sea | 27 |
| JFLOUNSETO | Japanese flounder | *Paralichthys olivaceus* | Seto Inland Sea | 20 |
| JFLOUNNSJ | Japanese flounder | *Paralichthys olivaceus* | Sea of Japan (north) | 15 |
| JFLOUNECS | Japanese flounder | *Paralichthys olivaceus* | Sea of Japan and East China Sea | 28 |
| RONOFLOUNSOJ | Round-nose flounder | *Eopsetta grigorjewi* | Sea of Japan | 21 |
| POFLOUNSOJ | Point-head flounder | *Cleisthenes herzensteini* | Sea of Japan | 17 |
| WILLFLOUNNP | Willowy flounder | *Tanakius kitaharae* | Pacific | 16 |
| JPPUFFSOJ | Japanese pufferfish | *Takifugu rubripes* | Sea of Japan, East China Sea, and Seto Inland Sea | 12 |
| JPPUFFISE | Japanese pufferfish | *Takifugu rubripes* | Ise and Mikawa Bay | 21 |

### Appendix S6. Results of MSE simulation using the stochastic HS SR curve.

Here we present the simulation results for Trajectory Scenarios L, M, and H.

When there is no error other than R variation, NEW is the best performer in terms of the last stage of management for Trajectory Scenario L (Table S6–1, and Figs. S6–1 and S6–2). However, when other uncertainties are added in the simulations, NEW2 generally outperforms NEW with more balanced results in performance statistics. While the conservation performance of NEW2 is comparable to that of 40–10, the early-stage management performance of NEW2 is superior to that of 40–10 (Table S6–1, Columns of CAT.F and CC.F). For Trajectory Scenario M (Table S6–2, and Figs. S6–3 and S6–4) and Trajectory Scenario H (Table S6–3, and Figs. S6–5 and S6–6), NEW2 also outperforms NEW especially when there are errors on total biomass and SB estimates. In general, 40–10 slightly outperforms NEW2 for Trajectory Scenario H except for AAV of catch in the last stage of management. 40–10 provides the worst AAV performance as a whole. From a comprehensive perspective, NEW2 is well-balanced and suitable for the management objectives in both of the early and last stages of management.

Table S6–1. Average performance measures for tested HCRs in Trajectory Scenario L ($SB_{msy} \rightarrow 0.2SB_{msy}$). SS = Sensitivity Scenarios.

| SS1 | | SB | CAT.L | AAV | CAT.F | CC.F |
|---|---|---|---|---|---|---|
| Status Quo | No time lag | 0.742 | 0.920 | 0.082 | 0.287 | 0.772 |
| | A time lag | 0.529 | 0.627 | 0.108 | 0.235 | 0.568 |
| 40-10 | No time lag | 1.474 | 0.954 | 0.102 | 0.126 | 0.290 |
| | A time lag | 1.480 | 0.969 | 0.118 | 0.079 | 0.152 |
| NEW | No time lag | 1.000 | 0.932 | 0.063 | 0.285 | 0.786 |
| | A time lag | 0.969 | 0.904 | 0.066 | 0.256 | 0.645 |
| NEW2 | No time lag | 1.314 | 0.937 | 0.050 | 0.256 | 0.705 |
| | A time lag | 1.330 | 0.949 | 0.052 | 0.219 | 0.546 |
| $F_{msy}^{TRUE}$ | No time lag | 0.909 | 0.909 | 0.046 | 0.293 | 0.810 |

| SS5 | | SB | CAT.L | AAV | CAT.F | CC.F |
|---|---|---|---|---|---|---|
| Status Quo | No time lag | 0.753 | 0.916 | 0.102 | 0.292 | 0.725 |
| | A time lag | 0.469 | 0.543 | 0.146 | 0.258 | 0.584 |
| 40-10 | No time lag | 1.480 | 0.953 | 0.107 | 0.133 | 0.258 |
| | A time lag | 1.479 | 0.953 | 0.135 | 0.093 | 0.166 |
| NEW | No time lag | 1.125 | 0.961 | 0.073 | 0.280 | 0.713 |
| | A time lag | 1.100 | 0.939 | 0.083 | 0.250 | 0.555 |
| NEW2 | No time lag | 1.431 | 0.942 | 0.063 | 0.242 | 0.607 |
| | A time lag | 1.444 | 0.949 | 0.067 | 0.212 | 0.456 |
| $F_{msy}^{TRUE}$ | No time lag | 0.923 | 0.923 | 0.057 | 0.295 | 0.803 |

| SS2 | | SB | CAT.L | AAV | CAT.F | CC.F |
|---|---|---|---|---|---|---|
| Status Quo | No time lag | 0.882 | 0.944 | 0.115 | 0.286 | 0.704 |
| | A time lag | 0.621 | 0.617 | 0.127 | 0.254 | 0.557 |
| 40-10 | No time lag | 1.400 | 0.943 | 0.123 | 0.167 | 0.342 |
| | A time lag | 1.440 | 0.964 | 0.135 | 0.119 | 0.223 |
| NEW | No time lag | 1.282 | 0.978 | 0.092 | 0.261 | 0.636 |
| | A time lag | 1.217 | 0.934 | 0.097 | 0.245 | 0.551 |
| NEW2 | No time lag | 1.531 | 0.917 | 0.078 | 0.231 | 0.570 |
| | A time lag | 1.536 | 0.921 | 0.082 | 0.204 | 0.450 |
| $F_{msy}^{TRUE}$ | No time lag | 0.937 | 0.937 | 0.077 | 0.298 | 0.807 |

| SS6 | | SB | CAT.L | AAV | CAT.F | CC.F |
|---|---|---|---|---|---|---|
| Status Quo | No time lag | 0.683 | 0.833 | 0.360 | 0.290 | 0.769 |
| | A time lag | 0.398 | 0.465 | 0.431 | 0.255 | 0.574 |
| 40-10 | No time lag | 1.468 | 0.965 | 0.540 | 0.155 | 0.323 |
| | A time lag | 1.415 | 0.928 | 0.510 | 0.104 | 0.181 |
| NEW | No time lag | 1.088 | 0.956 | 0.265 | 0.282 | 0.786 |
| | A time lag | 1.047 | 0.908 | 0.264 | 0.247 | 0.575 |
| NEW2 | No time lag | 1.388 | 0.934 | 0.228 | 0.249 | 0.670 |
| | A time lag | 1.403 | 0.938 | 0.221 | 0.214 | 0.504 |
| $F_{msy}^{TRUE}$ | No time lag | 0.866 | 0.886 | 0.261 | 0.296 | 0.873 |

| SS3 | | SB | CAT.L | AAV | CAT.F | CC.F |
|---|---|---|---|---|---|---|
| Status Quo | No time lag | 0.677 | 0.831 | 0.324 | 0.285 | 0.810 |
| | A time lag | 0.402 | 0.461 | 0.456 | 0.237 | 0.591 |
| 40-10 | No time lag | 1.439 | 0.955 | 0.585 | 0.143 | 0.340 |
| | A time lag | 1.414 | 0.954 | 0.536 | 0.085 | 0.169 |
| NEW | No time lag | 1.018 | 0.949 | 0.256 | 0.281 | 0.799 |
| | A time lag | 0.956 | 0.884 | 0.253 | 0.245 | 0.633 |
| NEW2 | No time lag | 1.354 | 0.956 | 0.231 | 0.252 | 0.707 |
| | A time lag | 1.328 | 0.938 | 0.218 | 0.211 | 0.537 |
| $F_{msy}^{TRUE}$ | No time lag | 0.837 | 0.859 | 0.261 | 0.292 | 0.832 |

| SS7 | | SB | CAT.L | AAV | CAT.F | CC.F |
|---|---|---|---|---|---|---|
| Status Quo | No time lag | 0.643 | 0.715 | 0.602 | 0.312 | 0.851 |
| | A time lag | 0.461 | 0.465 | 0.640 | 0.292 | 0.617 |
| 40-10 | No time lag | 1.474 | 0.973 | 0.731 | 0.211 | 0.450 |
| | A time lag | 1.360 | 0.886 | 0.755 | 0.189 | 0.349 |
| NEW | No time lag | 1.246 | 0.911 | 0.500 | 0.292 | 0.838 |
| | A time lag | 1.131 | 0.807 | 0.502 | 0.284 | 0.610 |
| NEW2 | No time lag | 1.506 | 0.875 | 0.456 | 0.256 | 0.662 |
| | A time lag | 1.501 | 0.870 | 0.445 | 0.234 | 0.509 |
| $F_{msy}^{TRUE}$ | No time lag | 0.742 | 0.813 | 0.500 | 0.307 | 0.990 |

| SS4 | | SB | CAT.L | AAV | CAT.F | CC.F |
|---|---|---|---|---|---|---|
| Status Quo | No time lag | 0.366 | 0.461 | 0.662 | 0.290 | 0.828 |
| | A time lag | 0.246 | 0.294 | 0.715 | 0.264 | 0.670 |
| 40-10 | No time lag | 1.265 | 0.915 | 0.990 | 0.185 | 0.474 |
| | A time lag | 1.066 | 0.788 | 0.919 | 0.117 | 0.242 |
| NEW | No time lag | 0.826 | 0.795 | 0.565 | 0.284 | 0.810 |
| | A time lag | 0.690 | 0.670 | 0.551 | 0.256 | 0.683 |
| NEW2 | No time lag | 1.217 | 0.917 | 0.465 | 0.260 | 0.743 |
| | A time lag | 1.171 | 0.865 | 0.438 | 0.232 | 0.599 |
| $F_{msy}^{TRUE}$ | No time lag | 0.619 | 0.682 | 0.499 | 0.297 | 0.848 |

Table S6–2. Average performance measures for tested HCRs in Trajectory Scenario M ($SB_{msy} \rightarrow SB_{msy}$). SS = Sensitivity Scenarios.

| SS1 | | SB | CAT.L | AAV | CAT.F | CC.F |
|---|---|---|---|---|---|---|
| Status Quo | No time lag | 1.172 | 0.984 | 0.062 | 0.940 | 0.945 |
| | A time lag | 1.084 | 0.904 | 0.070 | 0.921 | 0.922 |
| 40-10 | No time lag | 1.549 | 0.954 | 0.088 | 0.699 | 0.703 |
| | A time lag | 1.552 | 0.960 | 0.102 | 0.664 | 0.665 |
| NEW | No time lag | 1.264 | 0.983 | 0.052 | 0.930 | 0.937 |
| | A time lag | 1.225 | 0.975 | 0.055 | 0.920 | 0.923 |
| NEW2 | No time lag | 1.502 | 0.954 | 0.045 | 0.797 | 0.803 |
| | A time lag | 1.509 | 0.950 | 0.046 | 0.773 | 0.776 |
| $F_{msy}^{TRUE}$ | No time lag | 1.004 | 1.004 | 0.042 | 0.995 | 1.003 |

| SS5 | | SB | CAT.L | AAV | CAT.F | CC.F |
|---|---|---|---|---|---|---|
| Status Quo | No time lag | 1.254 | 0.997 | 0.079 | 0.908 | 0.911 |
| | A time lag | 1.128 | 0.900 | 0.094 | 0.891 | 0.890 |
| 40-10 | No time lag | 1.608 | 0.911 | 0.095 | 0.633 | 0.635 |
| | A time lag | 1.642 | 0.932 | 0.112 | 0.596 | 0.596 |
| NEW | No time lag | 1.442 | 0.950 | 0.064 | 0.866 | 0.878 |
| | A time lag | 1.435 | 0.958 | 0.070 | 0.859 | 0.865 |
| NEW2 | No time lag | 1.672 | 0.896 | 0.057 | 0.740 | 0.749 |
| | A time lag | 1.681 | 0.902 | 0.059 | 0.714 | 0.718 |
| $F_{msy}^{TRUE}$ | No time lag | 0.997 | 0.997 | 0.055 | 0.986 | 1.003 |

| SS2 | | SB | CAT.L | AAV | CAT.F | CC.F |
|---|---|---|---|---|---|---|
| Status Quo | No time lag | 1.524 | 0.910 | 0.086 | 0.803 | 0.814 |
| | A time lag | 1.432 | 0.854 | 0.094 | 0.791 | 0.795 |
| 40-10 | No time lag | 1.598 | 0.935 | 0.110 | 0.711 | 0.715 |
| | A time lag | 1.595 | 0.925 | 0.116 | 0.667 | 0.671 |
| NEW | No time lag | 1.511 | 0.922 | 0.081 | 0.821 | 0.840 |
| | A time lag | 1.489 | 0.937 | 0.085 | 0.808 | 0.824 |
| NEW2 | No time lag | 1.722 | 0.885 | 0.072 | 0.704 | 0.720 |
| | A time lag | 1.742 | 0.897 | 0.075 | 0.674 | 0.689 |
| $F_{msy}^{TRUE}$ | No time lag | 1.003 | 1.003 | 0.073 | 1.003 | 1.025 |

| SS6 | | SB | CAT.L | AAV | CAT.F | CC.F |
|---|---|---|---|---|---|---|
| Status Quo | No time lag | 1.179 | 0.983 | 0.335 | 0.961 | 0.979 |
| | A time lag | 0.977 | 0.822 | 0.353 | 0.941 | 0.966 |
| 40-10 | No time lag | 1.490 | 0.952 | 0.544 | 0.780 | 0.791 |
| | A time lag | 1.489 | 0.947 | 0.506 | 0.728 | 0.745 |
| NEW | No time lag | 1.219 | 0.956 | 0.259 | 0.988 | 1.010 |
| | A time lag | 1.156 | 0.913 | 0.258 | 0.970 | 0.999 |
| NEW2 | No time lag | 1.507 | 0.943 | 0.231 | 0.848 | 0.868 |
| | A time lag | 1.488 | 0.933 | 0.228 | 0.818 | 0.842 |
| $F_{msy}^{TRUE}$ | No time lag | 0.966 | 0.999 | 0.261 | 1.042 | 1.071 |

| SS3 | | SB | CAT.L | AAV | CAT.F | CC.F |
|---|---|---|---|---|---|---|
| Status Quo | No time lag | 1.151 | 0.990 | 0.316 | 0.957 | 0.961 |
| | A time lag | 0.990 | 0.846 | 0.329 | 0.929 | 0.936 |
| 40-10 | No time lag | 1.430 | 0.974 | 0.621 | 0.791 | 0.795 |
| | A time lag | 1.355 | 0.943 | 0.573 | 0.749 | 0.755 |
| NEW | No time lag | 1.002 | 0.945 | 0.276 | 1.028 | 1.033 |
| | A time lag | 0.878 | 0.822 | 0.266 | 1.017 | 1.025 |
| NEW2 | No time lag | 1.311 | 0.970 | 0.233 | 0.890 | 0.895 |
| | A time lag | 1.298 | 0.965 | 0.228 | 0.866 | 0.874 |
| $F_{msy}^{TRUE}$ | No time lag | 0.965 | 0.991 | 0.258 | 1.023 | 1.029 |

| SS7 | | SB | CAT.L | AAV | CAT.F | CC.F |
|---|---|---|---|---|---|---|
| Status Quo | No time lag | 1.459 | 0.897 | 0.554 | 0.877 | 0.941 |
| | A time lag | 1.185 | 0.713 | 0.573 | 0.831 | 0.901 |
| 40-10 | No time lag | 1.474 | 1.005 | 0.769 | 0.919 | 0.977 |
| | A time lag | 1.318 | 0.847 | 0.764 | 0.892 | 0.966 |
| NEW | No time lag | 1.236 | 0.892 | 0.513 | 1.018 | 1.087 |
| | A time lag | 1.007 | 0.722 | 0.512 | 1.011 | 1.089 |
| NEW2 | No time lag | 1.512 | 0.897 | 0.457 | 0.866 | 0.927 |
| | A time lag | 1.436 | 0.841 | 0.453 | 0.844 | 0.910 |
| $F_{msy}^{TRUE}$ | No time lag | 0.884 | 0.973 | 0.490 | 1.095 | 1.212 |

| SS4 | | SB | CAT.L | AAV | CAT.F | CC.F |
|---|---|---|---|---|---|---|
| Status Quo | No time lag | 0.898 | 0.850 | 0.570 | 0.976 | 0.983 |
| | A time lag | 0.734 | 0.673 | 0.599 | 0.973 | 0.982 |
| 40-10 | No time lag | 1.163 | 0.877 | 1.074 | 0.888 | 0.894 |
| | A time lag | 0.901 | 0.694 | 1.015 | 0.872 | 0.882 |
| NEW | No time lag | 0.670 | 0.694 | 0.620 | 1.108 | 1.114 |
| | A time lag | 0.498 | 0.519 | 0.616 | 1.128 | 1.140 |
| NEW2 | No time lag | 1.084 | 0.895 | 0.505 | 0.982 | 0.988 |
| | A time lag | 0.882 | 0.737 | 0.508 | 0.972 | 0.983 |
| $F_{msy}^{TRUE}$ | No time lag | 0.764 | 0.840 | 0.495 | 1.055 | 1.062 |

Table S6–3. Average performance measures for tested HCRs in Trajectory Scenario H ($SB_{msy} \rightarrow 1.8SB_{msy}$). SS = Sensitivity Scenarios.

| SS1 | | SB | CAT.L | AAV | CAT.F | CC.F |
|---|---|---|---|---|---|---|
| Status Quo | No time lag | 1.931 | 0.836 | 0.045 | 0.860 | 0.996 |
| | A time lag | 1.957 | 0.845 | 0.047 | 0.863 | 0.997 |
| 40-10 | No time lag | 1.869 | 0.877 | 0.066 | 0.926 | 1.099 |
| | A time lag | 1.859 | 0.889 | 0.071 | 0.930 | 1.104 |
| NEW | No time lag | 1.744 | 0.888 | 0.044 | 1.015 | 1.195 |
| | A time lag | 1.719 | 0.909 | 0.047 | 1.033 | 1.214 |
| NEW2 | No time lag | 1.975 | 0.841 | 0.039 | 0.857 | 1.008 |
| | A time lag | 1.986 | 0.843 | 0.041 | 0.856 | 1.005 |
| $F_{msy}^{TRUE}$ | No time lag | 1.012 | 1.012 | 0.041 | 1.346 | 1.595 |

| SS5 | | SB | CAT.L | AAV | CAT.F | CC.F |
|---|---|---|---|---|---|---|
| Status Quo | No time lag | 1.992 | 0.796 | 0.061 | 0.816 | 0.967 |
| | A time lag | 1.987 | 0.794 | 0.066 | 0.819 | 0.967 |
| 40-10 | No time lag | 2.053 | 0.810 | 0.072 | 0.844 | 1.042 |
| | A time lag | 2.063 | 0.798 | 0.080 | 0.847 | 1.047 |
| NEW | No time lag | 1.929 | 0.821 | 0.056 | 0.927 | 1.134 |
| | A time lag | 1.916 | 0.826 | 0.059 | 0.947 | 1.156 |
| NEW2 | No time lag | 2.197 | 0.758 | 0.050 | 0.782 | 0.955 |
| | A time lag | 2.206 | 0.750 | 0.053 | 0.787 | 0.959 |
| $F_{msy}^{TRUE}$ | No time lag | 0.991 | 0.991 | 0.054 | 1.355 | 1.674 |

| SS2 | | SB | CAT.L | AAV | CAT.F | CC.F |
|---|---|---|---|---|---|---|
| Status Quo | No time lag | 2.488 | 0.606 | 0.084 | 0.650 | 0.771 |
| | A time lag | 2.490 | 0.613 | 0.085 | 0.646 | 0.764 |
| 40-10 | No time lag | 2.131 | 0.784 | 0.077 | 0.800 | 1.002 |
| | A time lag | 2.080 | 0.777 | 0.080 | 0.796 | 1.003 |
| NEW | No time lag | 2.066 | 0.777 | 0.067 | 0.822 | 1.031 |
| | A time lag | 2.071 | 0.789 | 0.070 | 0.826 | 1.036 |
| NEW2 | No time lag | 2.301 | 0.723 | 0.062 | 0.687 | 0.862 |
| | A time lag | 2.299 | 0.725 | 0.063 | 0.683 | 0.856 |
| $F_{msy}^{TRUE}$ | No time lag | 1.006 | 1.006 | 0.072 | 1.370 | 1.738 |

| SS6 | | SB | CAT.L | AAV | CAT.F | CC.F |
|---|---|---|---|---|---|---|
| Status Quo | No time lag | 1.936 | 0.821 | 0.301 | 0.873 | 1.044 |
| | A time lag | 1.927 | 0.815 | 0.305 | 0.865 | 1.040 |
| 40-10 | No time lag | 1.792 | 0.878 | 0.423 | 1.000 | 1.247 |
| | A time lag | 1.822 | 0.873 | 0.406 | 1.017 | 1.277 |
| NEW | No time lag | 1.600 | 0.912 | 0.234 | 1.094 | 1.354 |
| | A time lag | 1.595 | 0.929 | 0.233 | 1.108 | 1.379 |
| NEW2 | No time lag | 1.912 | 0.846 | 0.222 | 0.922 | 1.138 |
| | A time lag | 1.928 | 0.848 | 0.213 | 0.924 | 1.150 |
| $F_{msy}^{TRUE}$ | No time lag | 0.971 | 0.996 | 0.267 | 1.406 | 1.745 |

| SS3 | | SB | CAT.L | AAV | CAT.F | CC.F |
|---|---|---|---|---|---|---|
| Status Quo | No time lag | 1.910 | 0.855 | 0.285 | 0.890 | 1.028 |
| | A time lag | 1.895 | 0.853 | 0.283 | 0.869 | 1.007 |
| 40-10 | No time lag | 1.639 | 0.929 | 0.495 | 1.057 | 1.255 |
| | A time lag | 1.619 | 0.930 | 0.469 | 1.070 | 1.279 |
| NEW | No time lag | 1.307 | 0.966 | 0.252 | 1.169 | 1.379 |
| | A time lag | 1.228 | 0.941 | 0.242 | 1.190 | 1.410 |
| NEW2 | No time lag | 1.719 | 0.912 | 0.218 | 0.988 | 1.164 |
| | A time lag | 1.689 | 0.910 | 0.214 | 0.999 | 1.181 |
| $F_{msy}^{TRUE}$ | No time lag | 0.965 | 0.996 | 0.258 | 1.375 | 1.624 |

| SS7 | | SB | CAT.L | AAV | CAT.F | CC.F |
|---|---|---|---|---|---|---|
| Status Quo | No time lag | 2.385 | 0.564 | 0.546 | 0.688 | 0.836 |
| | A time lag | 2.326 | 0.552 | 0.540 | 0.675 | 0.840 |
| 40-10 | No time lag | 1.655 | 0.918 | 0.687 | 1.174 | 1.565 |
| | A time lag | 1.622 | 0.889 | 0.676 | 1.174 | 1.600 |
| NEW | No time lag | 1.460 | 0.884 | 0.478 | 1.216 | 1.622 |
| | A time lag | 1.399 | 0.852 | 0.479 | 1.232 | 1.675 |
| NEW2 | No time lag | 1.832 | 0.892 | 0.431 | 1.019 | 1.352 |
| | A time lag | 1.750 | 0.843 | 0.425 | 1.027 | 1.399 |
| $F_{msy}^{TRUE}$ | No time lag | 0.876 | 0.972 | 0.493 | 1.521 | 2.019 |

| SS4 | | SB | CAT.L | AAV | CAT.F | CC.F |
|---|---|---|---|---|---|---|
| Status Quo | No time lag | 1.799 | 0.861 | 0.534 | 0.902 | 1.039 |
| | A time lag | 1.708 | 0.817 | 0.522 | 0.892 | 1.032 |
| 40-10 | No time lag | 1.220 | 0.916 | 1.052 | 1.238 | 1.480 |
| | A time lag | 0.966 | 0.748 | 1.015 | 1.281 | 1.542 |
| NEW | No time lag | 0.735 | 0.743 | 0.620 | 1.381 | 1.644 |
| | A time lag | 0.577 | 0.589 | 0.592 | 1.452 | 1.736 |
| NEW2 | No time lag | 1.142 | 0.913 | 0.495 | 1.195 | 1.418 |
| | A time lag | 1.055 | 0.837 | 0.484 | 1.237 | 1.473 |
| $F_{msy}^{TRUE}$ | No time lag | 0.766 | 0.846 | 0.500 | 1.426 | 1.686 |

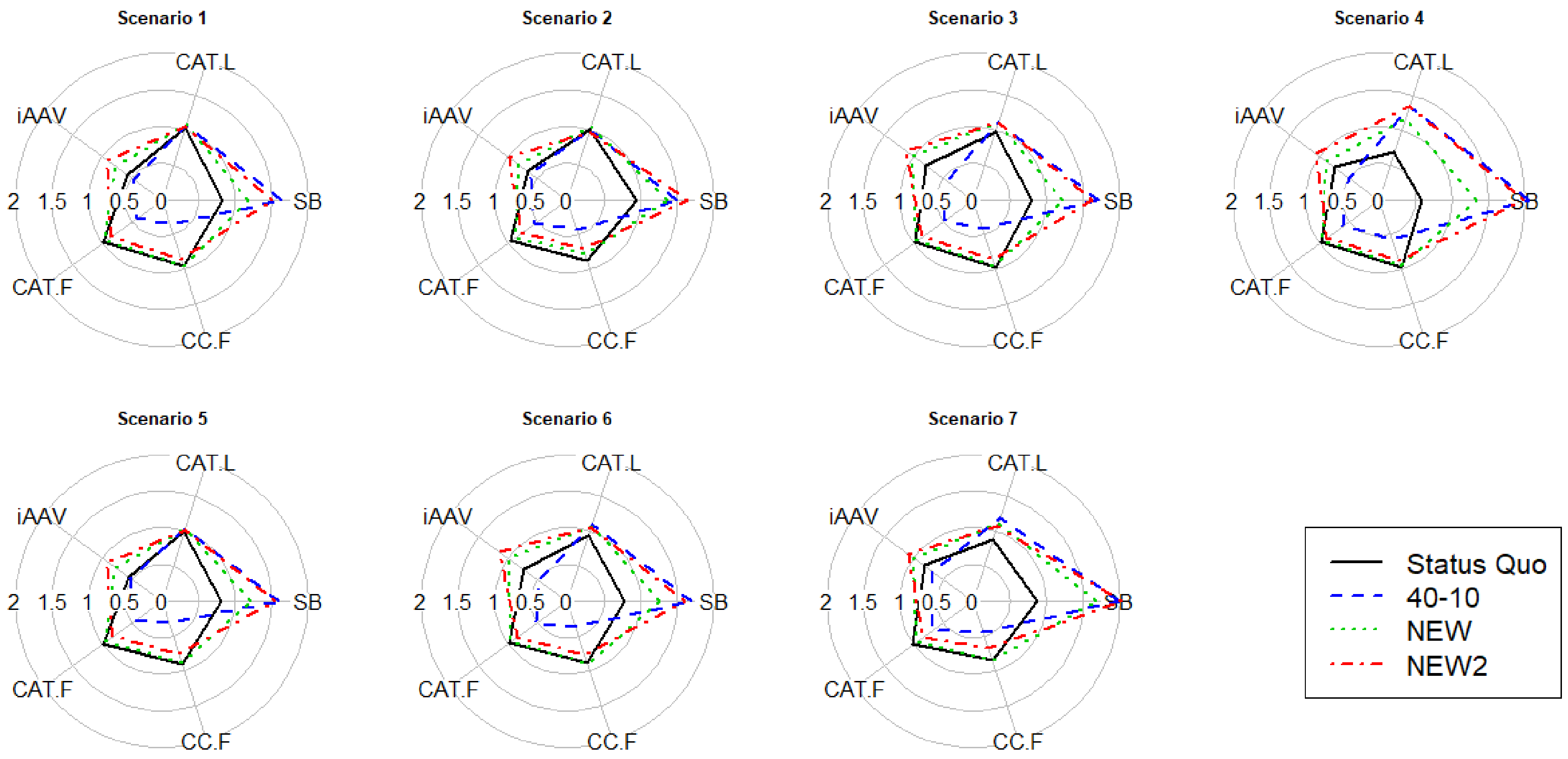


Fig. S6–1. The radar chart plots of performance measures for Trajectory Scenario L over Sensitivity Scenarios when there is no time lag. The explanation of performance measures was provided in Table 3. Status Quo is the traditional HCR in Japan, 40–10 is the 40–10 rule, NEW is the new HCR with $\beta$=1.0 and NEW2 is the new HCR with $\beta$=0.8. The circle with radius 1 denotes the results harvested by the true $F_{\mathrm{msy}}$.

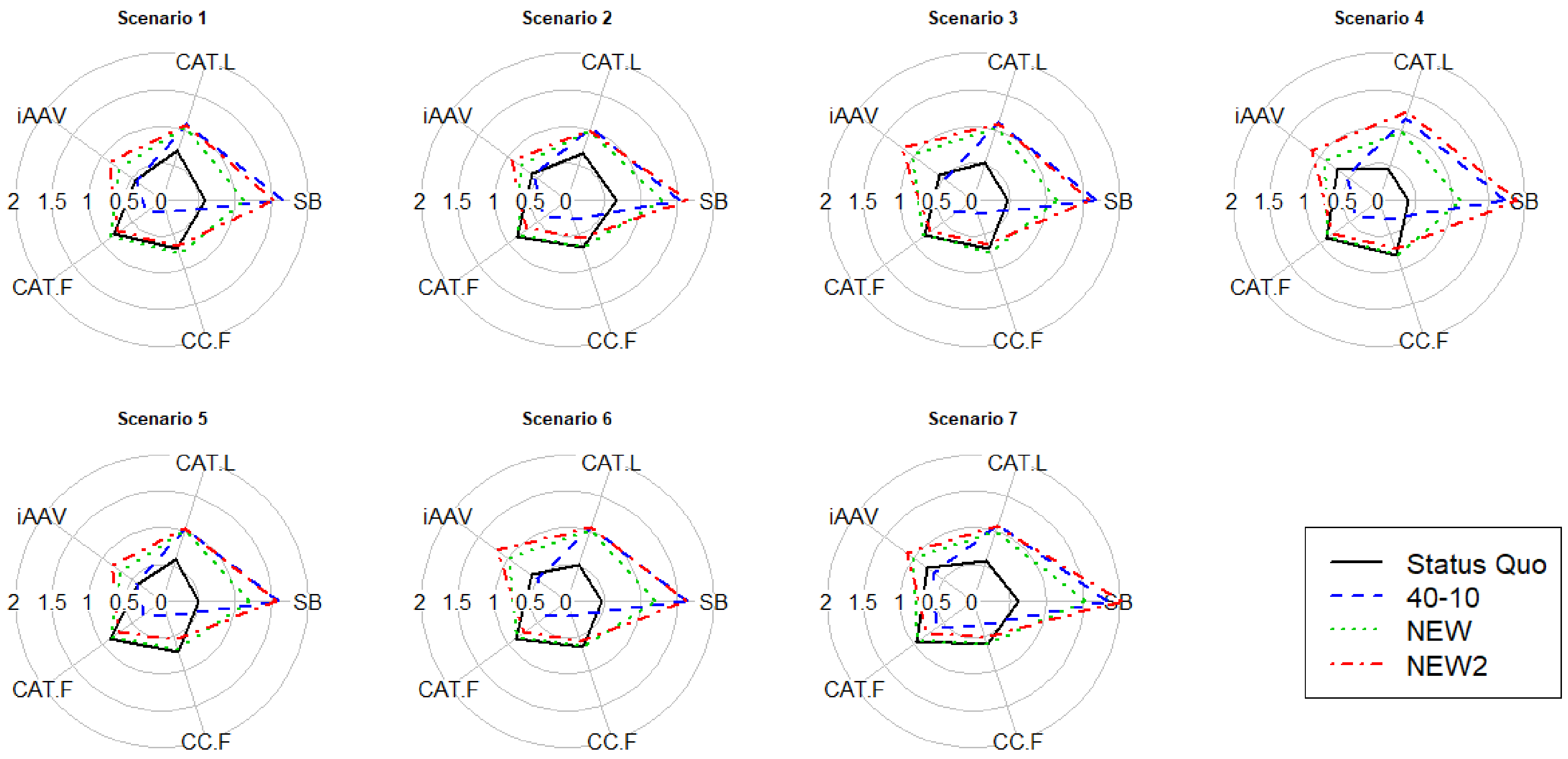


Fig. S6–2. The radar chart plots of performance measures for Trajectory Scenario L over Sensitivity Scenarios when there is a time lag. The explanation of performance measures was provided in Table 3. Status Quo is the traditional HCR in Japan, 40–10 is the 40–10 rule, NEW is the new HCR with $\beta$=1.0 and NEW2 is the new HCR with $\beta$=0.8. The circle with radius 1 denotes the results harvested by the true $F_{msy}$.

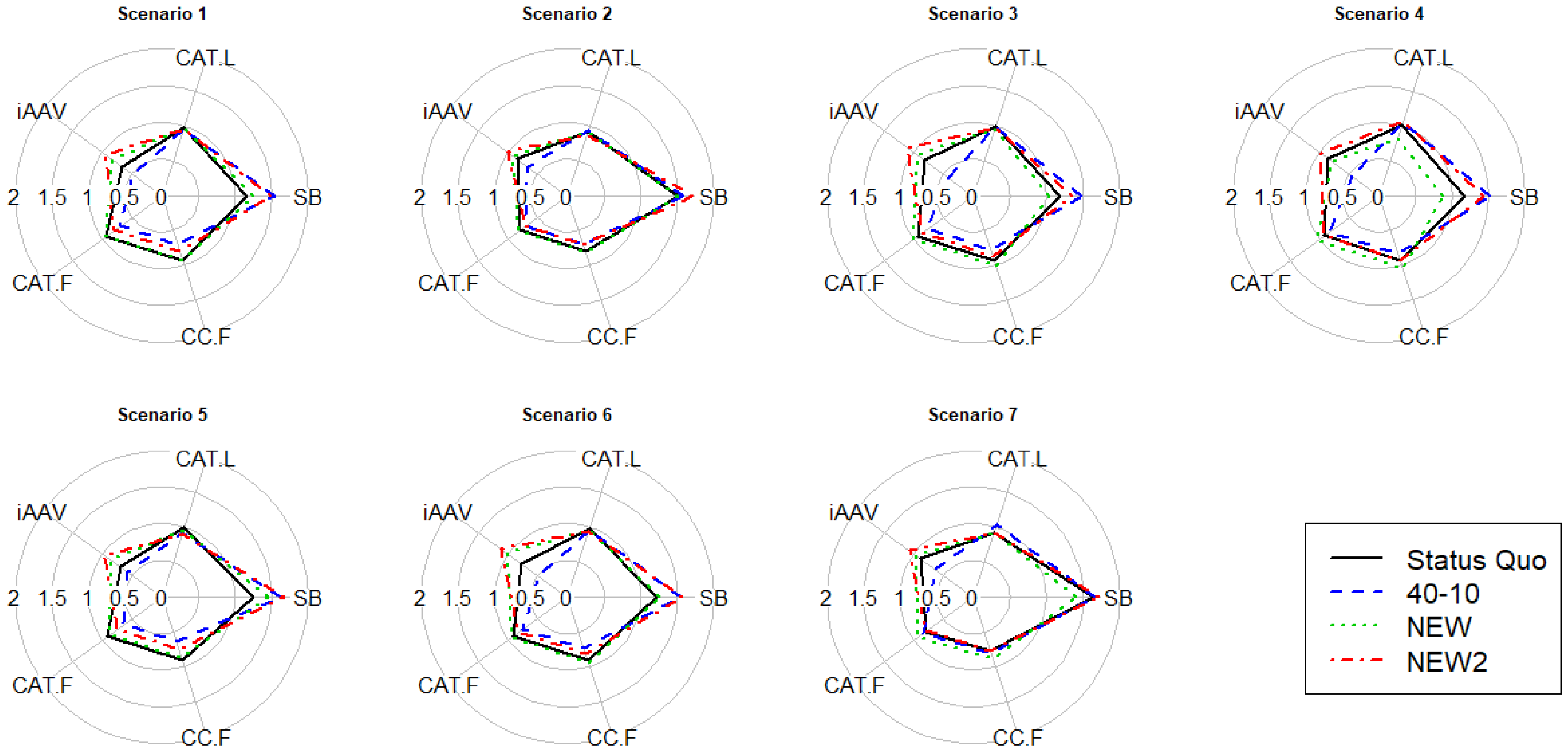


Fig. S6–3. The radar chart plots of performance measures for Trajectory Scenario M over Sensitivity Scenarios when there is no time lag. The explanation of performance measures was provided in Table 3. Status Quo is the traditional HCR in Japan, 40–10 is the 40–10 rule, NEW is the new HCR with $\beta$=1.0 and NEW2 is the new HCR with $\beta$=0.8. The circle with radius 1 denotes the results harvested by the true $F_{msy}$.

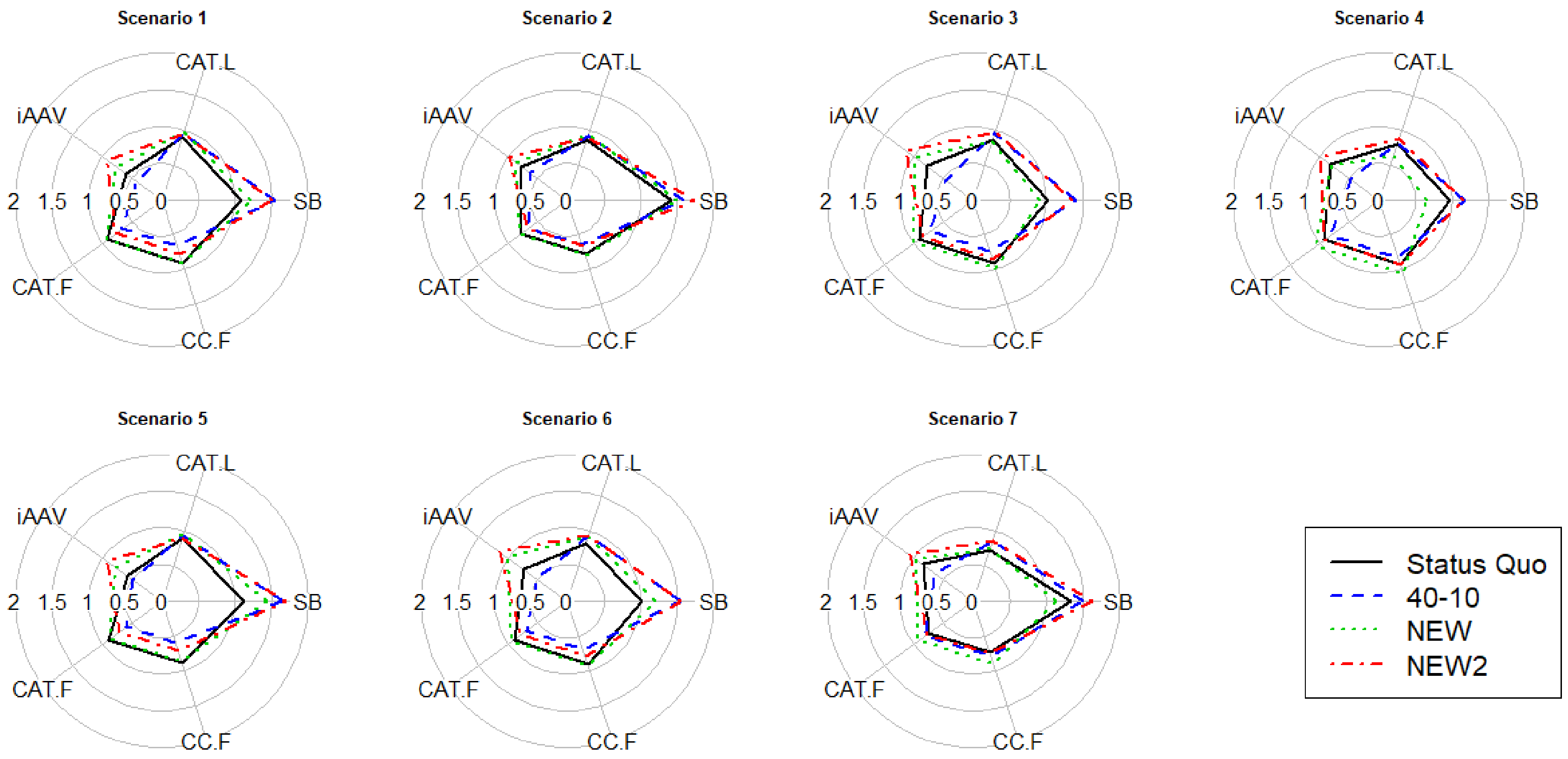


Fig. S6–4. The radar chart plots of performance measures for Trajectory Scenario M over Sensitivity Scenarios when there is a time lag. The explanation of performance measures was provided in Table 3. Status Quo is the traditional HCR in Japan, 40–10 is the 40–10 rule, NEW is the new HCR with $\beta$=1.0 and NEW2 is the new HCR with $\beta$=0.8. The circle with radius 1 denotes the results harvested by the true $F_{msy}$.

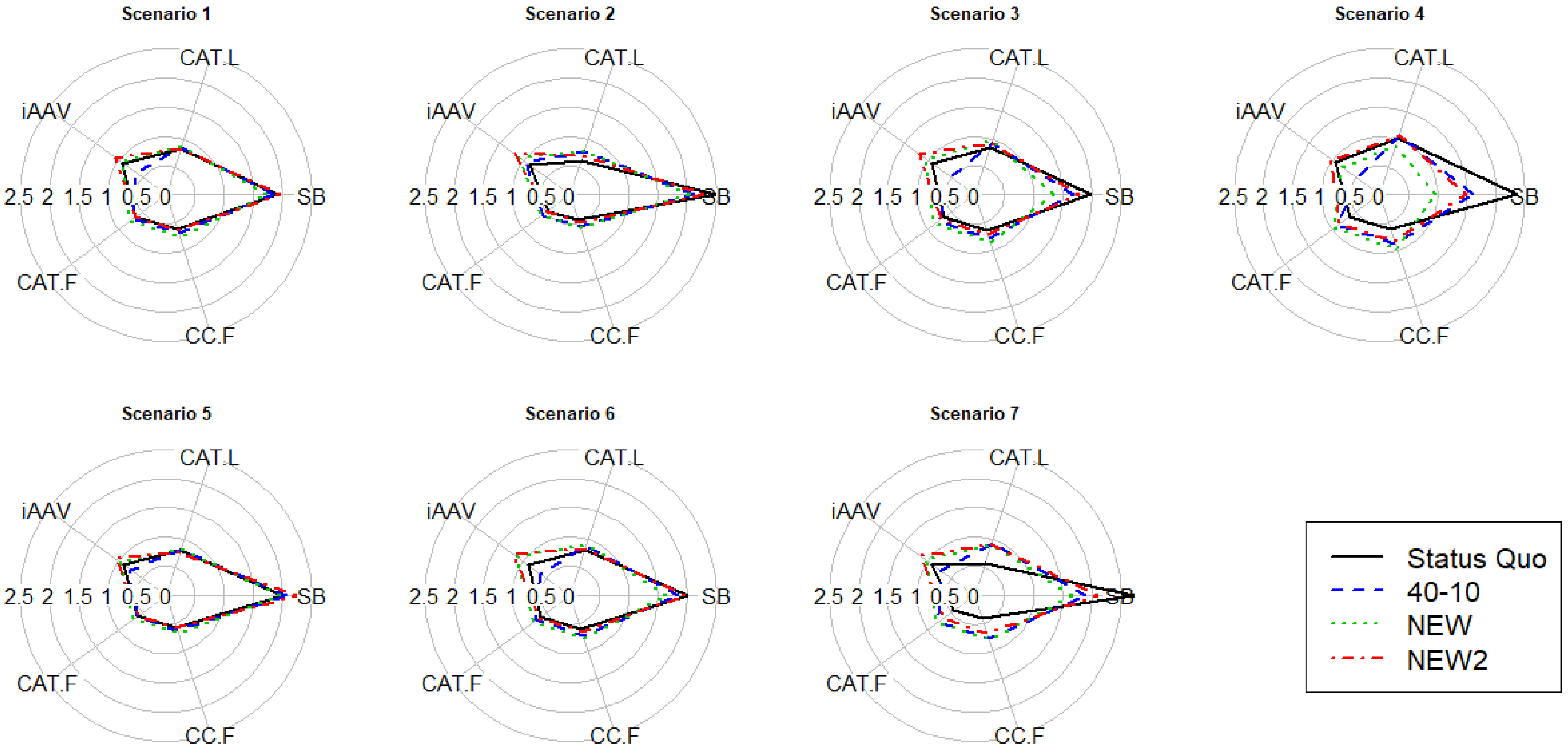


Fig. S6–5. The radar chart plots of performance measures for Trajectory Scenario H over Sensitivity Scenarios when there is no time lag. The explanation of performance measures was provided in Table 3. Status Quo is the traditional HCR in Japan, 40–10 is the 40–10 rule, NEW is the new HCR with $\beta$=1.0 and NEW2 is the new HCR with $\beta$=0.8. The circle with radius 1 denotes the results harvested by the true $F_{msy}$.

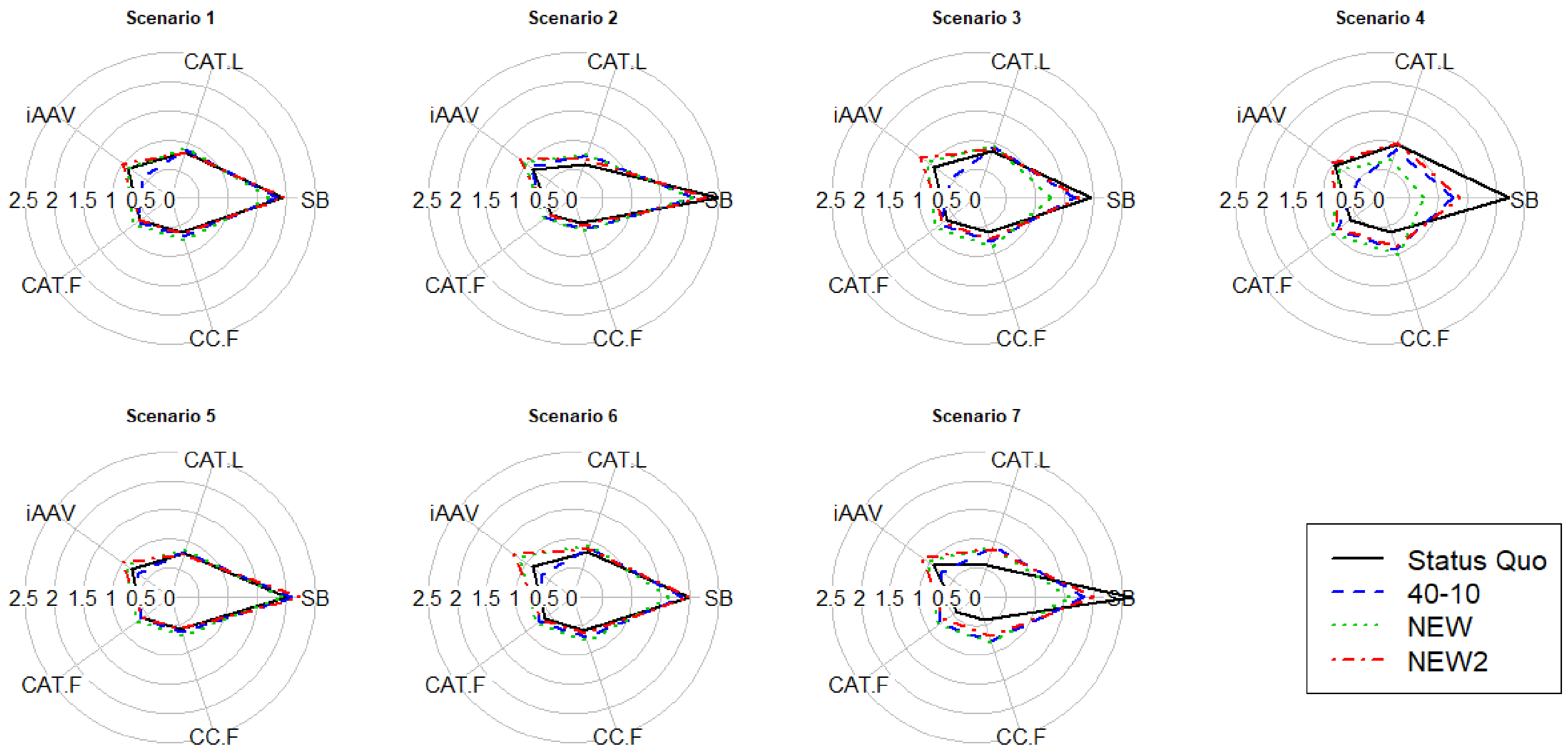


Fig. S6–6. The radar chart plots of performance measures for Trajectory Scenario H over Sensitivity Scenarios when there is a time lag. The explanation of performance measures was provided in Table 3. Status Quo is the traditional HCR in Japan, 40–10 is the 40–10 rule, NEW is the new HCR with $\beta$=1.0 and NEW2 is the new HCR with $\beta$=0.8. The circle with radius 1 denotes the results harvested by the true $F_{msy}$.

#### Appendix S7. Recovery time to $SB_{msy}$ for the Trajectory Scenario L

We investigate how the change of the $\beta$ parameter in the new HCR affects the recovery time to $SB_{msy}$ for Trajectory Scenario L. $\beta$ is changed from 1.0 to 0.7 by 0.1. The results are provided by weighting the Bio-parameter Scenarios by their plausibility probabilities separately for Sensitivity Scenarios 1 to 7 (Fig. S7–1). In general, the $\beta$ equal to or less than 0.8 is enough to attain about 50% probability that SB exceeds $SB_{msy}$ on average for most of Sensitivity Scenarios except Scenario 7 (the worst case).

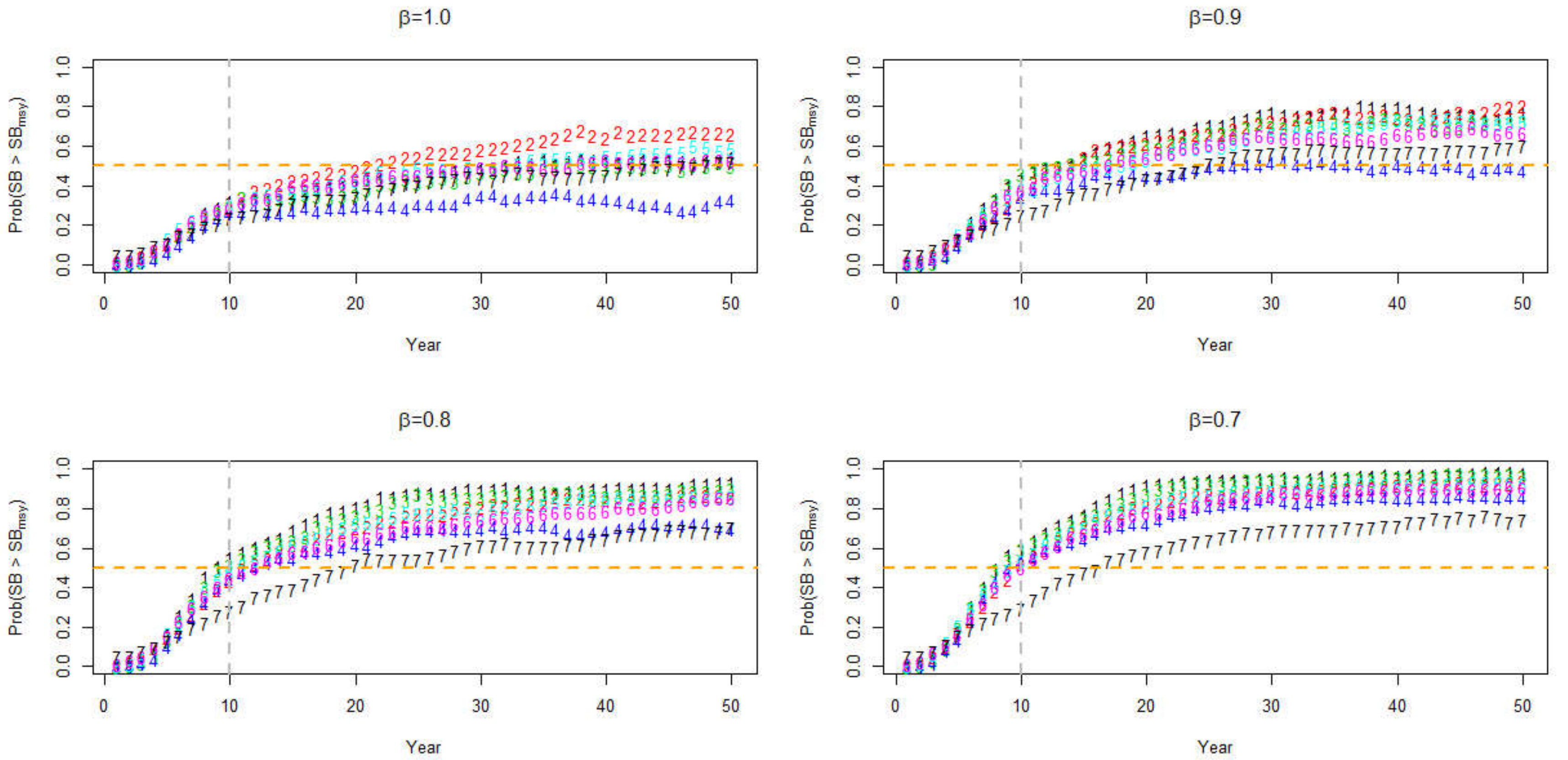


Fig. S7−1. The trajectories of SB over $SB_{msy}$ for Sensitivity Scenarios when there is a time lag. The horizontal broken line denotes the line at which Prob(SB > $SB_{msy}$)=0.5 and the horizontal broken line denote 10 years after the start of the new HCR management with $\beta$ = 1.0, 0.9, 0.8, and 0.7.

#### Appendix S8. Results of MSE simulation for Additional Sensitivity Scenarios.

Here we present the simulation results for Additional Sensitivity Scenarios.

When the errors of biomass estimates are autocorrelated, the performances of each HCR are slightly deteriorated (Table S6–1 (SS3) vs. Table S7–1 (ASS1)). Similarly, when there are the random errors in catch, the performances are mildly affected (Table S7–1 (ASS2)). When there is bias in biomass estimate, the resultant performance statistics are deteriorated especially for the true $F_{msy}$ (Table S7–1 (ASS3)). However, 40–10 and NEW2 work well for conservation purposes and NEW2 does not lose much catch at the early stage of management. When the Beverton-Holt SR curve is correct, the use of the HS SR curve incurs overfished SB by each HCR (Table S7–1 (ASS4)). However, the probability of population crash is generally small, and the catches for 40–10 and NEW2 are close to MSY. When the Ricker SR curve is correct, the use of the HS SR curve incurs underexploitation by each HCR (Table S7–1 (ASS5)). In the real situation, since the SR curve is reviewed every 5 years, we will not always continue to use the HS SR curve and therefore the probabilities of overexploitaion and uderexploitation would be overestimated by the simulation with continuously using the same SR curve as in this paper. For the Ricker SR curve scenario, 40–10 incurs a considerably small catch in the early and last stages of management (Table S7–1 (ASS5)). As a whole perspective, again, NEW2 is well-balanced and suitable for the management objectives in both of the early and last stages even for Additional Sensitivity Scenarios.

Table S8–1. Average performance measures for tested HCRs in Trajectory Scenario L ($SB_{msy} \rightarrow 0.2SB_{msy}$). ASS = Additional Sensitivity Scenarios.

| ASS1 | | SB | CAT.L | AAV | CAT.F | CC.F |
|---|---|---|---|---|---|---|
| Status Quo | No time lag | 0.646 | 0.773 | 0.379 | 0.274 | 0.807 |
| | A time lag | 0.399 | 0.452 | 0.471 | 0.225 | 0.587 |
| 40-10 | No time lag | 1.491 | 0.948 | 0.640 | 0.139 | 0.349 |
| | A time lag | 1.425 | 0.917 | 0.610 | 0.084 | 0.171 |
| NEW | No time lag | 1.046 | 0.937 | 0.297 | 0.272 | 0.811 |
| | A time lag | 0.940 | 0.840 | 0.290 | 0.238 | 0.652 |
| NEW2 | No time lag | 1.379 | 0.939 | 0.262 | 0.248 | 0.737 |
| | A time lag | 1.360 | 0.928 | 0.262 | 0.202 | 0.544 |
| $F_{msy}^{TRUE}$ | No time lag | 0.817 | 0.845 | 0.299 | 0.282 | 0.841 |

| ASS4 | | SB | CAT.L | AAV | CAT.F | CC.F |
|---|---|---|---|---|---|---|
| Status Quo | No time lag | 0.406 | 0.770 | 0.061 | 0.424 | 0.789 |
| | A time lag | 0.369 | 0.681 | 0.073 | 0.383 | 0.698 |
| 40-10 | No time lag | 0.729 | 0.951 | 0.086 | 0.245 | 0.445 |
| | A time lag | 0.736 | 0.956 | 0.095 | 0.181 | 0.319 |
| NEW | No time lag | 0.517 | 0.843 | 0.052 | 0.438 | 0.815 |
| | A time lag | 0.520 | 0.851 | 0.055 | 0.412 | 0.753 |
| NEW2 | No time lag | 0.743 | 0.932 | 0.045 | 0.385 | 0.716 |
| | A time lag | 0.745 | 0.937 | 0.046 | 0.354 | 0.647 |
| $F_{msy}^{TRUE}$ | No time lag | 0.985 | 0.985 | 0.029 | 0.331 | 0.621 |

| ASS2 | | SB | CAT.L | AAV | CAT.F | CC.F |
|---|---|---|---|---|---|---|
| Status Quo | No time lag | 0.685 | 0.844 | 0.101 | 0.278 | 0.754 |
| | A time lag | 0.428 | 0.497 | 0.129 | 0.224 | 0.551 |
| 40-10 | No time lag | 1.488 | 0.970 | 0.094 | 0.122 | 0.286 |
| | A time lag | 1.469 | 0.966 | 0.115 | 0.068 | 0.134 |
| NEW | No time lag | 1.048 | 0.956 | 0.067 | 0.271 | 0.766 |
| | A time lag | 0.981 | 0.891 | 0.071 | 0.241 | 0.619 |
| NEW2 | No time lag | 1.375 | 0.958 | 0.052 | 0.250 | 0.695 |
| | A time lag | 1.362 | 0.947 | 0.053 | 0.200 | 0.511 |
| $F_{msy}^{TRUE}$ | No time lag | 0.856 | 0.862 | 0.057 | 0.281 | 0.783 |

| ASS5 | | SB | CAT.L | AAV | CAT.F | CC.F |
|---|---|---|---|---|---|---|
| Status Quo | No time lag | 0.028 | 0.040 | 0.169 | 0.386 | 0.892 |
| | A time lag | 0.014 | 0.021 | 0.324 | 0.414 | 0.878 |
| 40-10 | No time lag | 2.597 | 0.182 | 0.060 | 0.071 | 0.169 |
| | A time lag | 2.891 | 0.184 | 0.065 | 0.047 | 0.102 |
| NEW | No time lag | 1.871 | 0.250 | 0.044 | 0.245 | 0.536 |
| | A time lag | 2.033 | 0.241 | 0.046 | 0.226 | 0.463 |
| NEW2 | No time lag | 2.128 | 0.213 | 0.042 | 0.214 | 0.465 |
| | A time lag | 2.454 | 0.213 | 0.043 | 0.189 | 0.388 |
| $F_{msy}^{TRUE}$ | No time lag | 0.754 | 0.754 | 0.253 | 0.589 | 1.311 |

| ASS3 | | SB | CAT.L | AAV | CAT.F | CC.F |
|---|---|---|---|---|---|---|
| Status Quo | No time lag | 0.615 | 0.770 | 0.090 | 0.281 | 0.778 |
| | A time lag | 0.408 | 0.420 | 0.114 | 0.242 | 0.592 |
| 40-10 | No time lag | 1.113 | 0.979 | 0.077 | 0.206 | 0.499 |
| | A time lag | 0.996 | 0.879 | 0.088 | 0.139 | 0.289 |
| NEW | No time lag | 0.860 | 0.925 | 0.066 | 0.283 | 0.793 |
| | A time lag | 0.704 | 0.724 | 0.074 | 0.262 | 0.645 |
| NEW2 | No time lag | 1.177 | 0.946 | 0.051 | 0.267 | 0.727 |
| | A time lag | 1.132 | 0.913 | 0.057 | 0.231 | 0.571 |
| $F_{msy}^{TRUE}$ | No time lag | 0.097 | 0.166 | 0.074 | 0.300 | 0.818 |

#### Appendix S9. Results of forecast and hindcast simulations using the real stock assessment outputs.

Here we present the forecast and hindcast simulation results using the real stock assessment outputs.

The time trajectories of $SB/SB_{msy}$ and $F/F_{msy}$ for the forecasting simulations are shown in Figs. S9–1 and S9–2. From a comprehensive standpoint, Status Quo fails preventing overfishing and recovering overfished populations. 40–10 and NEW2 can generally achieve the long-term management objectives. NEW has intermediate performances between 40–10 and NEW2. 40–10 needs a substantial catch reduction immediately after the introduction of HCR.

For JMACKTSST, Status Quo makes the stock extinct on average. Although F looks close to $F_{msy}$, this is because potentially high $F$s are excluded from averaging since $F$ becomes zero when the stock goes extinct (Fig. S9–2). Status Quo tends to provide $F$ higher than $F_{msy}$ ($F = 1.53 \times F_{msy}$ on average before extinction after the introduction of HCR). In addition to relatively high recruitment uncertainty ($CV_R = 0.47$, $\rho = 0.28$) and the time lag effect, such a high F leads the stock to extinction. NEW provides almost the same $F$ as $F_{msy}$ ($F = 0.98 \times F_{msy}$). Average $F$ for NEW2 is $0.80 \times F_{msy}$ and average $F$ for 40–10 is $0.74 \times F_{msy}$. Average catches for the last 10 years are 0.28×MSY for Status Quo, 0.94×MSY for 40–10, 0.97×MSY for NEW, and 0.96×MSY for NEW2. The similar things happen for CMACKTSST and SMACKPJPN so that Status Quo tends to cause extinction. For JPPUFFISE, Status Quo and NEW cause extinction with the 97% and 35% probabilities, respectively, while 40–10 and NEW2 have zero probabilities for extinction.

For JFLOUNNSJ, all HCRs make the stock recover. Average Fs of Status Quo and

NEW are close to $F_{msy}$, but the recovery time to $SB_{msy}$ is not enough for both HCRs. On the other hand, 40–10 and NEW2 permit the stock to recover to $SB_{msy}$ on average within 30 years after the introduction of HCR management. Increase of SB bring increase of catch and as a result, the average catches for the last 10 years are 0.34×MSY for Status Quo, 0.96×MSY for 40–10, 0.96×MSY for NEW, and 0.93×MSY for NEW2.

The time trajectories of $SB/SB_{msy}$ and $F/F_{msy}$ for the hindcasting simulations are shown in Figs. S9–3 and S9–4. The actually observed fishing rates are overall higher than $F_{msy}$ (Fig. S9–4). In most cases, SBs decreased or remained the low levels (Fig. S9–3). By contrast, NEW and NEW2 lead to stock recovery by reducing fishing pressures.

For APOLLPSOJ, NEW and NEW2 show a gradual increase of SB, whereas the observed SB remains a low level. By contrast, future projections do not show a conspicuous difference among HCRs and future SBs remain a very low level (Fig. S9–1). This is because the observed SB becomes a very low level at the observed final year and a very small catch is permitted by any HCR in future due to the delayed recovery (Neubauer et al. 2013). This suggests that the stock would have recovered if we had reduced catch before 10 years when SB was much higher than now.

We dealt with just 26 stocks that have VPA-based stock assessment results in this paper. However, there are more stocks assessed in Japan that use stock assessment models other than VPA or use different methods for data-limited species. Extending the simulations to those stocks will provide more exact pictures of fisheries stock status in Japan.

In sum, the simulations using the assessment results for data rich stocks in Japan indicate that management using a proper HCR leads to recovery of depleted stocks and increase of catches.

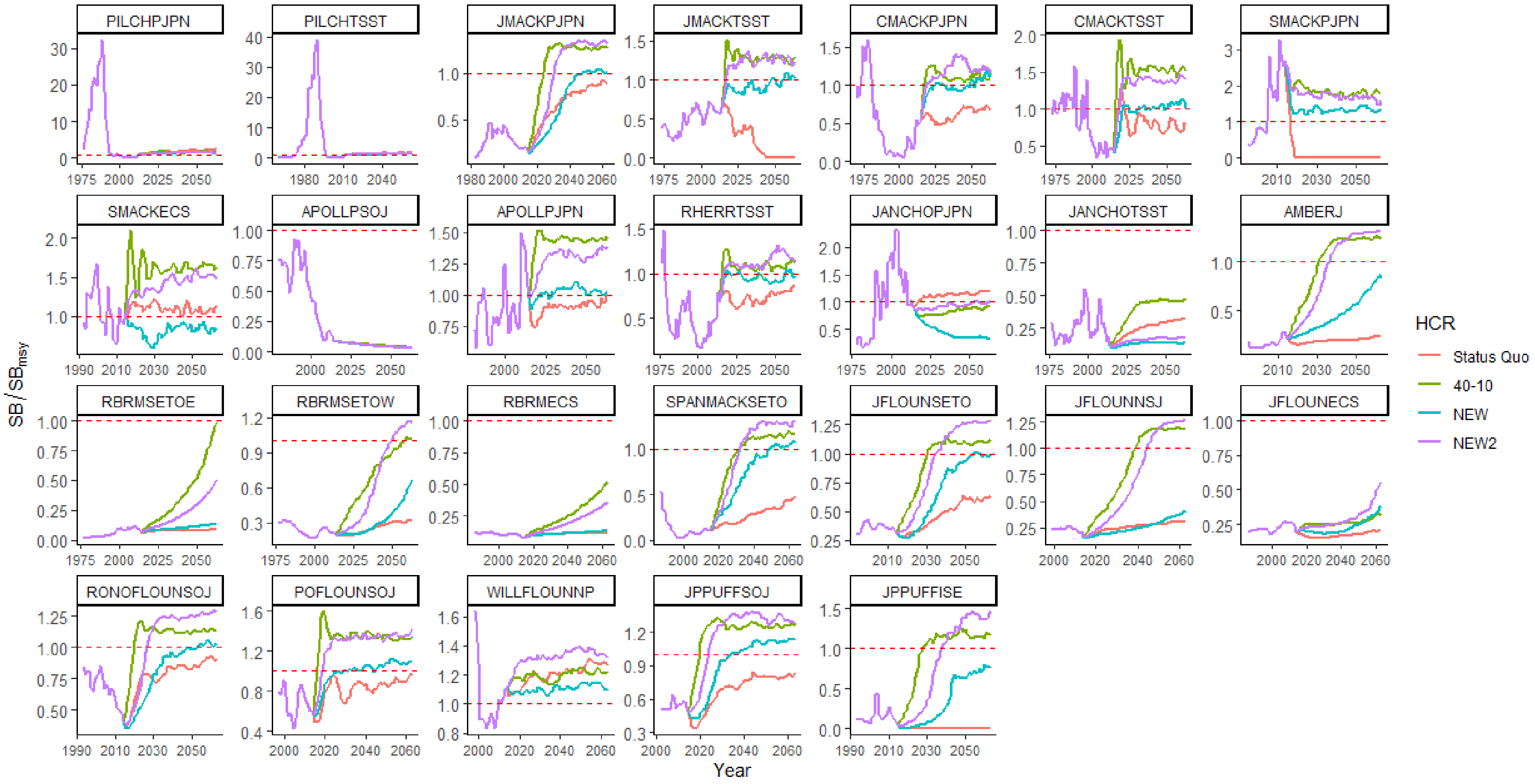


Fig. S9–1. The trajectories of spawning stock biomass from the forecast simulations for 26 Japanese data-rich stocks using the status quo, 40–10, NEW, and NEW2 HCRs. The broken red lines show $SB_{msy}$.

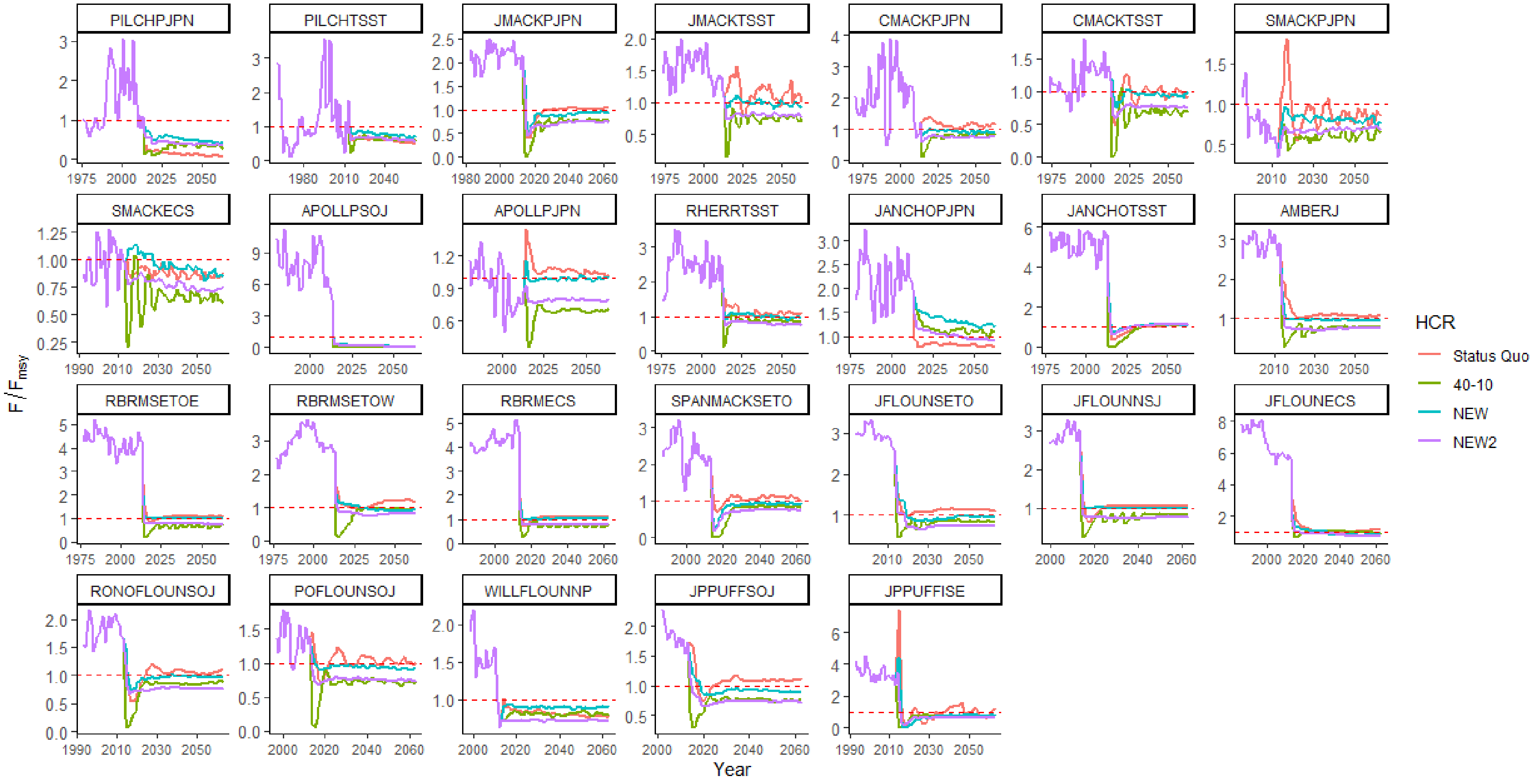


Fig. S9–2. The trajectories of fishing rate from the forecast simulations for 26 Japanese data-rich stocks using the status quo, 40–10, NEW, and NEW2 HCRs. The broken red lines show $SB_{msy}$.

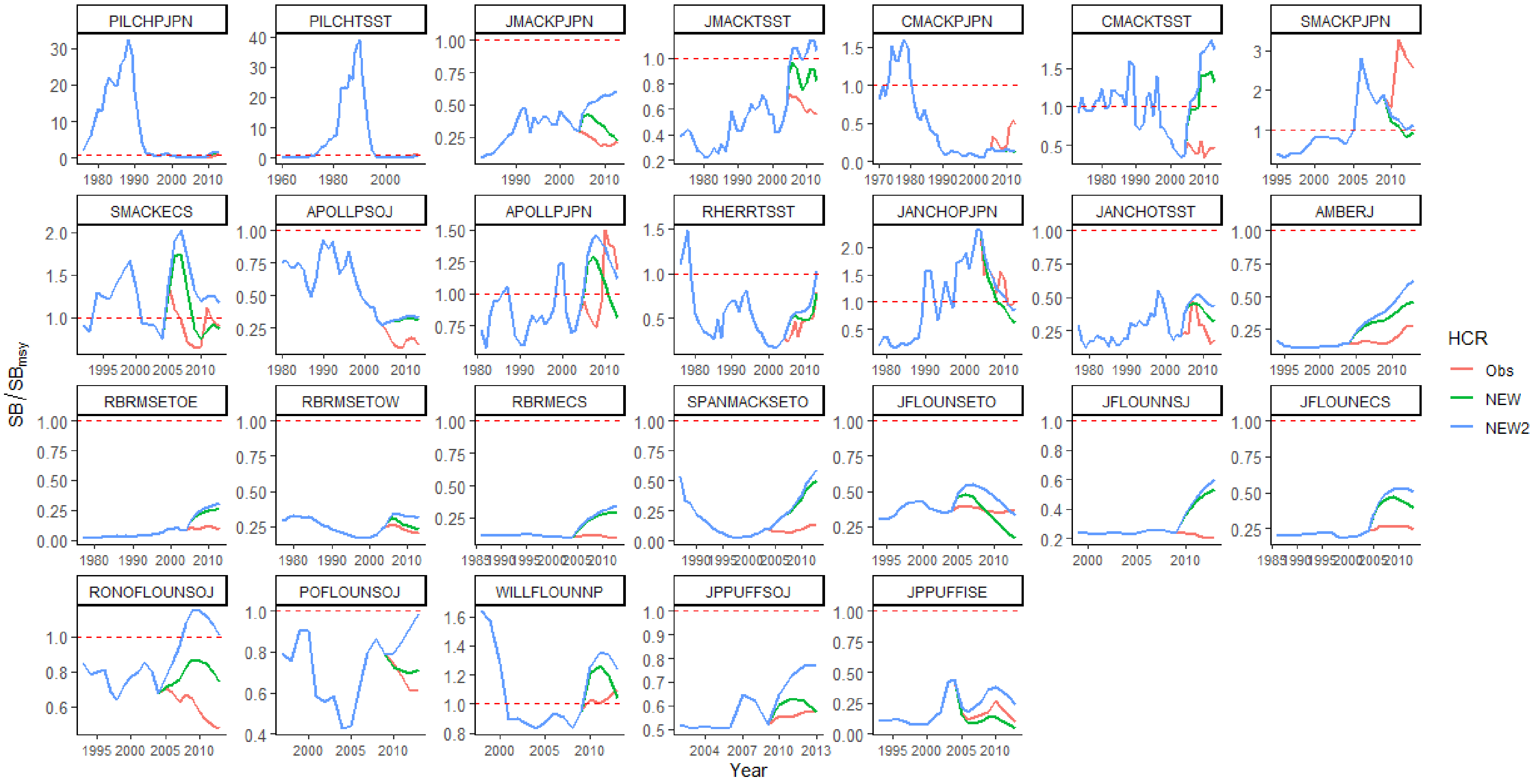


Fig. S9–3. The trajectories of spawning stock biomass from the hindcast simulations for 26 Japanese data-rich stocks. Obs denotes the actual observed spawning biomass. The broken red lines show $SB_{msy}$.

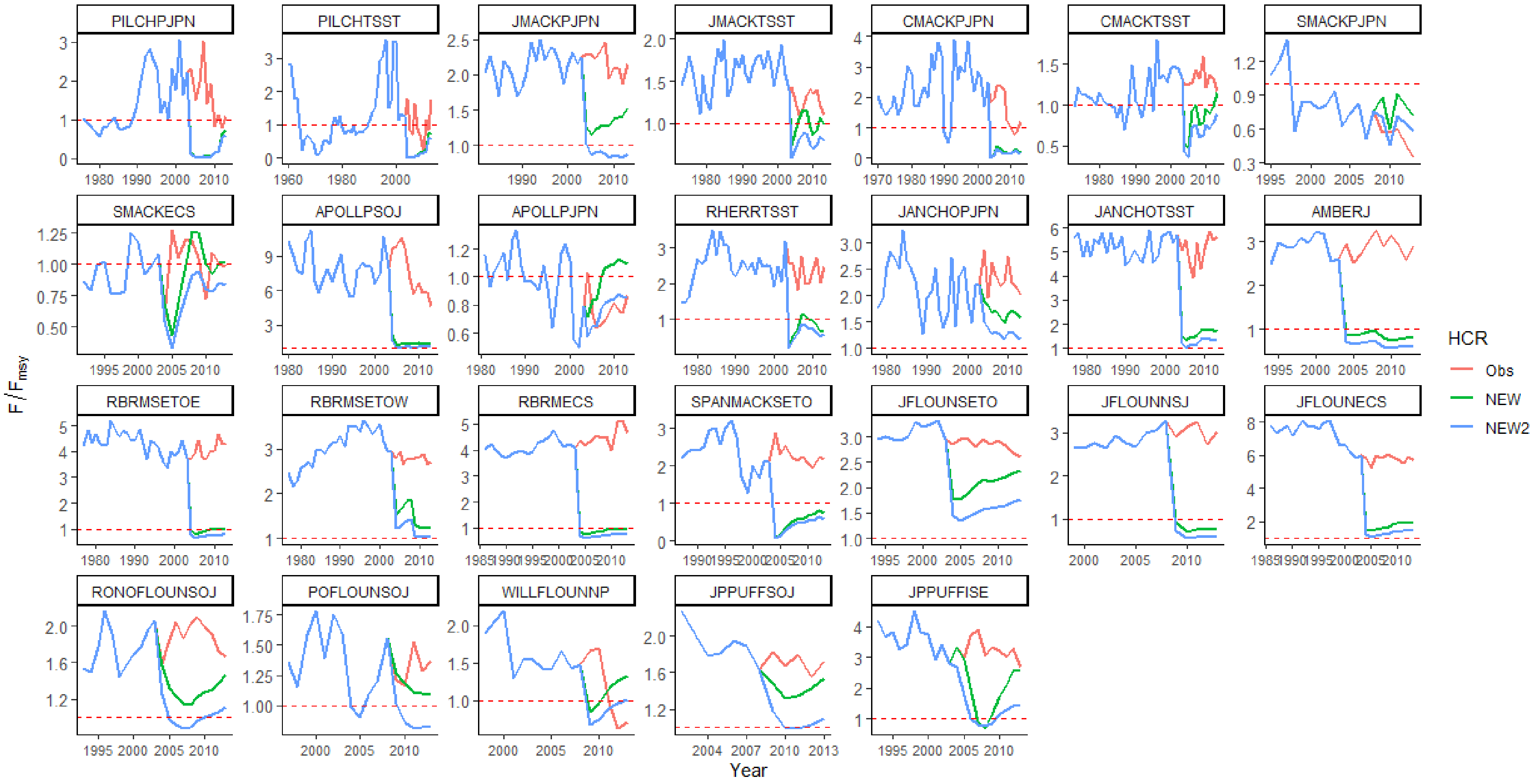


Fig. S9–4. The trajectories of fishing rate from the hindcast simulations for 26 Japanese data-rich stocks. Obs denotes the actual observed fishing rate. The broken red lines show $SB_{msy}$.
